# Group size modulates kinship dynamics and selection on social traits

**DOI:** 10.1101/2023.12.22.573047

**Authors:** Peng He, Michael N. Weiss, Samuel Ellis, Daniel W. Franks, Michael A. Cant, Darren P. Croft, Rufus A. Johnstone

**Author notes:** Correspondence (PH), (RAJ).

## Abstract

An individual’s relatedness to its group may change with age due to demographic processes, and such kinship dynamics can shape age-linked social behaviors. Existing models, however, have focused almost exclusively on dispersal and mating, leaving the consequences of group size local variation unexplored. Here, we extend these models to incorporate such variation within a single, genetically connected meta-population. We then contrast predicted kinship dynamics and age-linked selection for helping/harming in coexisting smaller and larger groups in three typical social mammal systems characterized by distinct patterns of dispersal and mating, followed by exploring how group size local variation impacts female and male kinship dynamics across broader demographic contexts to clarify the population-genetic mechanisms. We show that, within the adult reproductive lifespans of these social systems, an individual’s age-specific relatedness to others is higher, and changes faster (especially when younger), in smaller groups; consequently, smaller groups favor more extreme helping/harming (especially when younger) — in social system with bi-sexual philopatry with non-local mating (e.g., whales) that favors female shifts from harming to helping with age (which explains the evolution of menopause and post-reproductive helping), such shifts are earlier in smaller groups. Further explorations suggest that, in a genetically connected population with group size local variation, group size effects on local relatedness are shaped by the relative strengths of ancestry dilution versus lineage coalescence, and kinship dynamics in a group reflect not only its local demographic conditions, but also those in coexisting groups of different size. Collectively, our study generates new insights into how female and male kinship dynamics emerge and vary under within-population group-size heterogeneity, and how such locally dynamic, varying kinship environments may help to explain variation in age-linked trends in social traits, such as the timing of menopause in social mammals.

## INTRODUCTION

There is clear evidence that the mean relatedness of an individual to others in its social group may change systematically with age (Croft et al., 2021; Ellis et al., 2022). Such kinship dynamics, which are sensitive to a species’ demography, are predicted to influence sex- and age-specific social behaviors such as helping and harming (Croft et al., 2021; Ellis et al., 2022; Johnstone & Cant, 2008, 2010). For example, among social mammals, only humans and some toothed whales (Ellis et al., 2018) exhibit a life history pattern of early reproductive cessation (compared to males), followed by an extended period of post-reproductive life during which they assist kin (Croft et al., 2017; Johnstone & Cant, 2008, 2010, 2019). This unusual life history pattern is associated with unusual kinship dynamics: females in these species typically experience an increase in their relatedness to others in their social groups as they age, a trend that has been invoked to help explain the evolution of menopause and post-reproductive helping (Croft et al., 2017; Ellis et al., 2022; Johnstone & Cant, 2008, 2010).

Despite growing interest in kinship dynamics, existing models have focused chiefly on how these dynamics are shaped by typical rates of female and male philopatry versus dispersal (Cant & Johnstone, 2008; Johnstone & Cant, 2008), and of local versus non-local mating (Ellis et al., 2022; Johnstone & Cant, 2010). It remains unknown how local variation in demographic factors, such as social group size, modulates local kinship dynamics following population asymmetric subdivision, and the sex- and age-linked trends in social behaviors expected under such locally dynamic yet varying kinship environments. There is abundant evidence that group size varies within natural populations (Alexander, 1974; Brown & Brown, 1996; Brown et al., 2016; Crockett & Eisenberg, 1987; Dornhaus et al., 2012; Dunbar et al., 2018; Krause & Ruxton, 2002; Lehmann et al., 2007; Majolo et al., 2008; Safran, 2004), and that it is often correlated with social behaviors such as the amount of assistance provided by non-breeding helpers in cooperative breeders (Clutton-Brock et al., 2004; Farabaugh et al., 1992; Garcia-Ruiz & Taborsky, 2022; Josi et al., 2021; Pike et al., 2019). But the impacts of group size local variation on kinship dynamics and predicted changes in social behavior with sex and age remain unknown.

Here, we extend existing models of kinship dynamics (Ellis et al., 2022; Johnstone & Cant, 2010) to incorporate group size local deterministic variation within a genetically connected meta-population. We then use it to examine individuals’ age-specific relatedness to their groupmates in coexisting smaller and larger groups across the adult reproductive life stages, and explore how such emerging patterns of kinship dynamics influence age-linked selection for helping/harming (with a focus on social mammals), followed by discussing key challenges and uncertainties involved in empirically testing predictions on these age-linked social behaviors (using the killer whales *Orcinus orca* as an example). Finally, we explore our model in a broader range of demographic contexts to clarify the mechanisms through which demographic conditions, including group size local variation, influences emergent patterns of kinship dynamics females and males experience as they age. Together, our analyses provide new insights into how female and male kinship dynamics emerge and vary in coexisting local groups with deterministic differences in their sizes, and demonstrates a group-size-related mechanism that might be invoked to explain widespread variations in social traits in animal populations, such as the timing of reproductive cessation in social mammals (Nielsen et al., 2021; Thomas et al., 2001).

## METHODS

### Population demographic assumptions

Building on the baseline model of Johnstone and Cant (2010) and Ellis et al. (2022), we consider an infinite, diploid, bisexual population, divided into discrete social groups (a.k.a. islands, demes or patches, Rousset, 2004; Wright, 1931), a proportion *u* of which are of *α*-type consisting of *n*_*f⍺*_female and *n*_*m⍺*_ male breeders (i.e., adults who are reproductively active), while the remaining fraction 1 − *u* are of *β*-type consisting of *n*_*fβ*_ female and *n*_*mβ*_ male breeders (the baseline model assumes all groups have the same number of breeders). That is, grouptype is defined solely by the total number of breeders in a group, with *n*_*f⍺*_ + *n*_*m⍺*_ < *n*_*fβ*_ + *n*_*mβ*_, such that *⍺*-type groups are persistently smaller while *β*-type groups are persistently larger (when *n*_*f⍺*_ = *n*_*fβ*_ and *n*_*m⍺*_ = *n*_*mβ*_ it reduces to the baseline model representing a group-size homogeneous population). Such differences reflect deterministic (or structural) variation in the fixed numbers of breeders across coexisting grouptypes in a population, rather than their stochastic fluctuations (Rousset & Ronce, 2004) or statistical distributions. Also note that under the infinite island population model, inter-individual relatedness is generated within local groups (as it is consistently zero between groups).

As in the baseline model, time proceeds in discrete steps, in each of which every female produces some very large number, *p*, of offspring (regardless of group size), a fraction *x* of which are females while the rest are males (fecundity and primary sex ratio do not influence predictions but are introduced for clarity — see appendix for details; the assumption of a very large number of offspring in each sex makes sure that available breeding vacancies in each group at each timestep are always fully claimed by successful offspring — see below, such that group sizes remain consistent over time). At each timestep, the probability that an offspring born in a group is sired by a male from the same group is *m* in the baseline model (i.e., a measure of a local male’s mating success — with probability 1 − *m* the offspring is sired by a non-local male). That is, all the females and males considered in each group are breeders who reproduce at each timestep. Therefore, the model concerns the reproductive life history stages. In the context of group size local variation, this local siring probability is conditioned on grouptype in which the offspring is born, to account for differences in male numbers (see appendix for derivations). After reproduction, each female or male offspring disperses from its natal group (to a random group in the population) with probability *d*_*f*_ or *d*_*m*_, respectively. Then, offspring in each group (both those born locally that did not disperse, together with immigrants from the rest of the population) compete in a fair lottery manner for the available breeding vacancies created by the deaths in their own sex in that group — at each timestep each female or male breeder dies with probability *μ*_*f*_ or *μ*_*m*_, respectively. That is, mortality is modelled as constant sex-specific breeder turnover, approximating reproductive life history stages in which survival is relatively stable, such as adult lifespans after maturity and before late-life survival declines become pronounced in large mammals (Gaillard et al., 1998; Langtimm et al., 1998). In the presence of group size local variation, the probability that a native offspring wins a breeding vacancy (i.e., being recruited to its natal group a new breeder) is also conditioned on grouptype in which it is born (see derivations in the appendix). Offspring that fail to obtain any breeding vacancy die, and the cycle then repeats.

### Modelling kinship dynamics

Given above demographic assumptions, we derive (in the appendix) expressions for the expected relatedness of a breeder — with given sex and age (time since recruitment into the breeding pool), to a randomly sampled groupmate — with given sex but implicit age, distinguishing between smaller (*α*-type) and larger (*β*-type) groups. In the results section, we explore the predicted patterns of an individual’s sex-age-specific average relatedness to its groupmates in populations in which the proportion of *α*-type groups is either low (*u* = 0.1), medium (*u* = 0.5) or high (*u* = 0.9) — when *u* = 0 or *u* = 1 group-size heterogeneity disappears and our model reduces to the baseline model in which all groups are uniformly larger or smaller, respectively. For simplicity, we assume that *n*_*f⍺*_ = *n*_*m⍺*_ = 5 and *n*_*fβ*_ = *n*_*mβ*_ = 20 throughout (i.e., a smaller group contains 10 individuals, while a larger group contains 40 individuals, with the same number of males and females in either grouptype). The mortality rate of female and male breeders at each timestep across their lifespan in both grouptypes is *μ*_*f*_ = *μ*_*m*_ = 0.1. To make our explorations biologically relevant, we consider the three typical dispersal and mating patterns of social mammals addressed by Johnstone and Cant (2010) and Ellis et al. (2022), represented by (I) whales: *bisexual philopatry with non-local mating*, where *d*_*f*_ = 0.15, *d*_*m*_ = 0.15, and *m* = 0, (II) apes: *female-biased dispersal with local mating*, where *d*_*f*_ = 0.85, *d*_*m*_ = 0.15, and *m* = .82, and (III) typical mammals: *male-biased dispersal with local mating*, where *d*_*f*_ = 0.15, *d*_*m*_ = 0.85, and *m* = .82. To reveal the effects of group size variation on kinship dynamics, under each of these social mammal scenarios, we also show in the appendix for comparison the kinship dynamics in distinct, group-size homogeneous populations consisting of either all *α*-type or all *β*-type groups — as defined in the group-size ‘heterogeneous’ population, while keeping all other parameter values consistent. Besides, we also explore our model under other demographic contexts (including group size) to clarify how female and male kinship dynamics emerge and vary under group size local variation (see appendix for details).

### Evaluating selective pressures

We use the derived sex-age-explicit relatedness to quantify the strength of selection on helping/harming in both grouptypes, using the neighbor-modulated inclusive fitness approach (Ellis et al., 2022; Taylor & Frank, 1996). We assume helping/harming is genetically determined by a mutant allele, and the analysis proceeds in three steps (full expressions are given in the appendix). We first derive sex-grouptype-specific reproductive values under our demographic assumptions; then, we construct fitness functions for a focal individual in each sex-grouptype class, incorporating its own fecundity and mortality as well as those of others; next, we combine these fitness functions with the sex-age-explicit relatedness to examine the inclusive-fitness effect of the behavior expressed in given age, sex, and grouptype, provided it alters the fecundity or mortality of the focal individual and its groupmates (see fully details in the appendix).

More specifically, for the inclusive fitness calculations, we assume the behavior confers a total marginal fecundity or mortality benefits (or costs) of magnitude *b* (say 0 < |*b*| ≪ 1, where 0 > *b* for helping while *b* < 0 for harming) on the local recipients (either sex), distributed equally among them, and confers a constant, marginal fecundity or mortality cost *c* on the actor (0 < *c* ≪ 1; see details in the appendix). Then, the critical *b*^∗^ at which the net inclusive fitness effect is zero can be derived, and we then use the 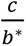 as a measure of sex-age-specific selective pressure on the behavior — the larger the positive value of 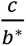, the stronger the selection for helping, whereas the smaller the negative value of 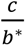, the stronger the selection for harming. To reveal how the prevalence of smaller groups in a population modulates selection, we explore populations with either low (*u* = 0.1), medium (*u* = 0.5) or high (*u* = 0.9) proportions of smaller groups. For comparison, we also show (in the appendix) the corresponding selective pressures in the counterpart, group-size homogeneous populations (i.e., as in Ellis et al., 2022) consisting entirely of smaller or entirely of larger groups (all other parameter values held constant).

## RESULTS

### Local variation in kinship dynamics

Across the whale, typical mammal, and the ape cases characterized by distinct patterns of female and male dispersal and prevalence of local mating, individuals’ average relatedness to their group is generally higher in smaller than larger groups. Of greater interest is the prediction that the rate of change in relatedness with age is typically higher in smaller than larger social groups — and that such differences are more pronounced at younger than older ages (Fig. 1). Across these cases, intra-population variation in group size does not alter the general, qualitative patterns of kinship dynamics experienced by individuals of either sex (Ellis et al., 2022), nor does the prevalence of smaller groups in the population. With given prevalence of smaller groups in the population, the differences in the patterns of kinship dynamics experienced by both sexes in smaller versus larger groups are most pronounced when both sexes are philopatric and mate non-locally (i.e., the whale case). With given patterns of mating and dispersal, the more prevalent the smaller groups are in the population (i.e., a higher *u*), the larger the differences in the patterns of kinship dynamics experienced by both sexes in smaller versus larger groups (Fig. 1). The predicted patterns and variations of female and male kinship dynamics under broader demographic contexts (i.e., varying magnitude of group size contrast, sex ratio, mortality, the prevalence of local mating, dispersal, and the proportion of smaller groups in the population) are enclosed in the appendix (Fig. A4-A7).

**Fig. 1.**
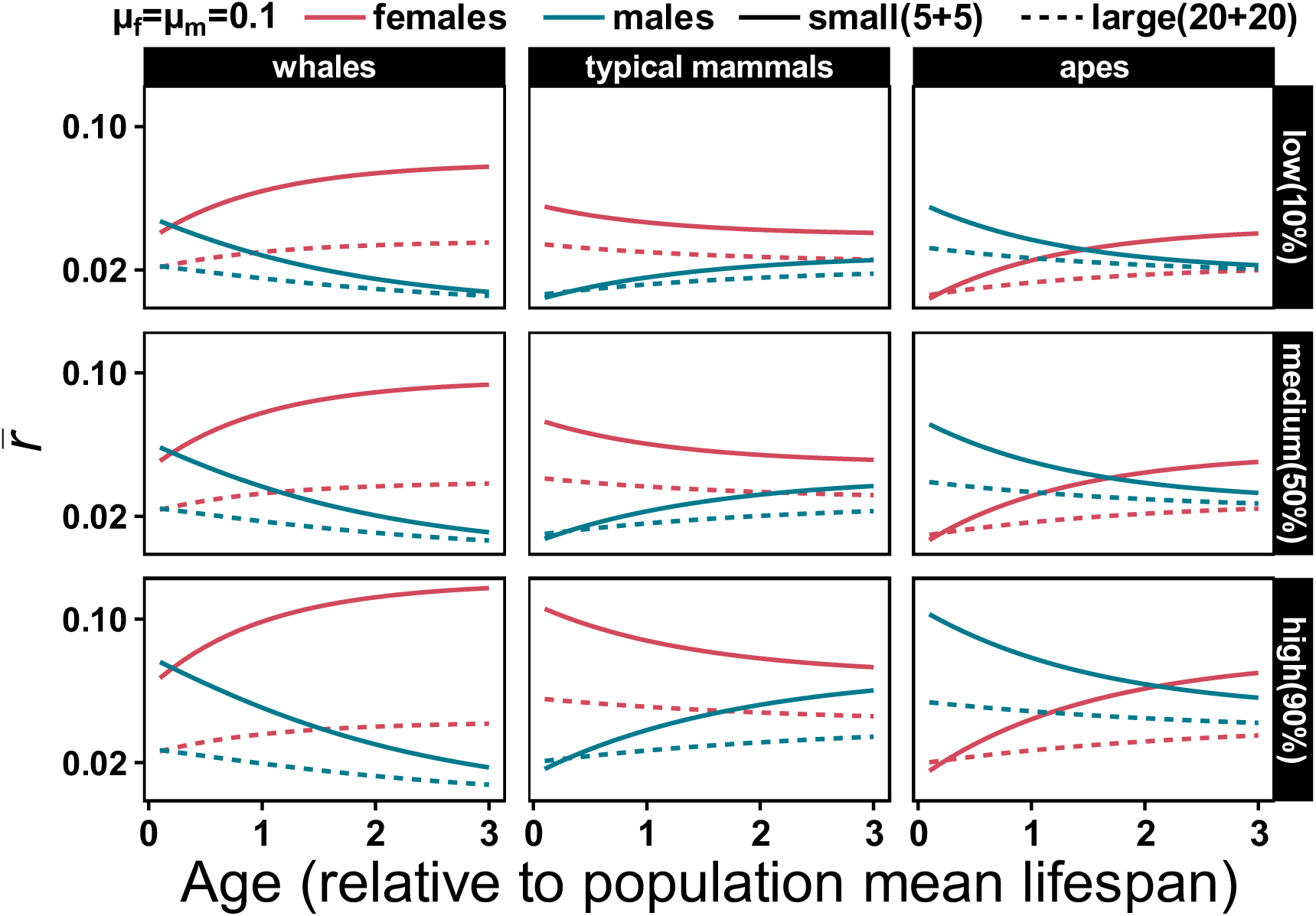
Predicted mean relatedness of an individual female (red) or male(blue) to others in its social group (smaller: 5 females and 5 males, solid lines; larger: 20 females and 20 males, dashed lines), as a function of its age (scaled relative to the mean lifespan under constant mortality *μ*_*f*_ = *μ*_*m*_ = 0.1 at each timestep). The columns in (A) or (B) show the three typical patterns of dispersal and mating that characterize social mammal systems (Ellis et al., 2022; Johnstone & Cant, 2010), referred to as: the ‘whale’ case (*bisexual philopatry with non-local mating*), the typical mammal case (*male-biased dispersal with local mating*), and the ‘ape’ case (*female-biased dispersal with local mating*), as detailed in methods. The rows show three different proportions of smaller groups in the population: low (*u* = 10%), medium (*u* = 50%), or high (*u* = 90%).

### Local variation in selection

We find that, across the whale, typical mammal, and the ape cases, selection for helping/harming is generally higher in smaller than larger groups, and the rate of change in selection with age is higher in smaller than larger social groups, especially at earlier than later life history stages (Fig. 2). Across these social mammal contexts, the presence of group size variation does not qualitatively alter the predicted patterns of selective pressures on the social behavior (Ellis et al., 2022), neither does the prevalence (*u*) of smaller groups in the population. In the whale and ape cases, we find that females in both smaller and larger groups experience a significant shift in selection — from being more harmful when younger to being more helpful when older, when the behavior affects fecundity, and that such shifts occur earlier in smaller groups (Fig. 2-A). With given prevalence of smaller groups in the population, the differences in the critical ages of these shifts between small and large groups are less significant in the ape than in the whale case; however, the impacts of the prevalence of smaller groups differs between the cases: in the whale case, such differences in critical age are more marked when smaller groups are more prevalent, while in the ape case such differences are most pronounced at low but least pronounced at intermediate prevalence of smaller groups in the population (Fig. 2-B).

**Fig. 2.**
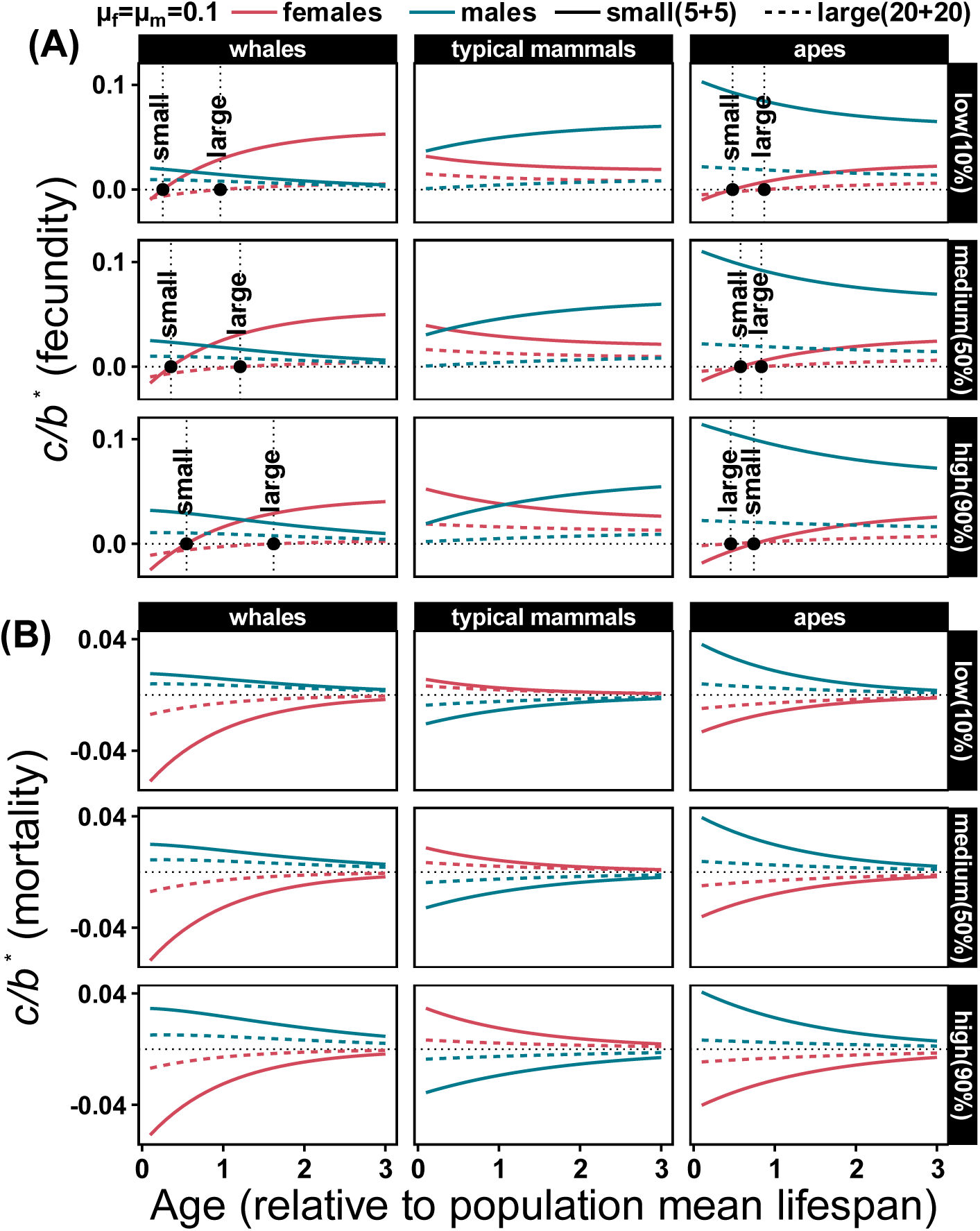
Predicted strength of selection for helping (positive values) or harming (negative values) as a function of sex (female: red; male: blue), age (scaled relative to the mean lifespan under constant mortality *μ*_*f*_ = *μ*_*m*_ = 0.1 at each timestep), and grouptype (solid lines: smaller groups; dashed lines: larger groups), given predicted patterns of sex-grouptype-specific kinship dynamics, provided the behavior affect the fecundity (A) or mortality (B) of an actor and the recipients. Dotted horizontal lines mark selectively neutral condition (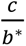 = 0) and dotted vertical lines in (A) mark the critical age (dots at curve intersections) at which selection on females shifts from favoring more harmful to favoring more helpful in smaller (5 females and 5 males) and larger (20 females and 20 males) groups. The columns in (A) or (B) show the three typical patterns of dispersal and mating that characterize social mammal systems (Ellis et al., 2022; Johnstone & Cant, 2010), referred to as: the ‘whale’ case *bisexual philopatry with non-local mating*), the typical mammal case (*male-biased dispersal with local mating* , and the ‘ape’ case *female-biased dispersal with local mating*), as detailed in methods. The rows show three different proportions of smaller groups in the population: low (*u* = 10%), medium (*u* = 50%), or high (*u* = 90%).

## DISCUSSION

Group size is a key component of social evolution (Bose et al., 2022; Brown & Brown, 1996; Brown et al., 2016; Dornhaus et al., 2012; Kaufhold & van Leeuwen, 2019; Krause & Ruxton, 2002; Majolo et al., 2008; Peña & Nöldeke, 2018; Powers & Lehmann, 2017). Building on existing models, we generated new insights into how patterns of female and male kinship dynamics emerge and vary in local groups following population asymmetric subdivision, and subsequently, how selection for age-linked social traits varies in such locally dynamic kinship environments.

Comparing distinct, group-size homogeneous populations across the whale, typical mammal, and ape contexts shows the consistency (Fig. A1) that individuals are more related to others in populations characterized by smaller groups (see also Dyble & Clutton-Brock, 2020; Powers & Lehmann, 2017). Our results across these social systems show that this expectation of higher relatedness in smaller social mammal groups extends to group-size heterogeneous populations (so does the expectation of stronger selection for helping/harming in smaller groups, Fig. A2). However, we argue that this correspondence is conditional rather than universal: the strength and even direction of local group size effects depend on sex, life history stage (age), breeder sex ratio, and social systems collectively realized by patterns of dispersal and local mating. For example, in the whale and ape contexts, an older male’s average relatedness to its groupmates becomes increasingly insensitive to group size (as differences between smaller and larger groups diminish with age); under unbiased, moderate dispersal and mortality between the sexes, combined with moderate local mating, males even exhibit consistently higher relatedness in larger groups (Fig. A4), a pattern contrary to the seemingly general expectation of higher relatedness in smaller social mammal groups.

More fundamentally, we find that an individual’s kinship ties to others within a local group reflects not only its own size, but also the demographic conditions experienced by the other grouptype (e.g., altering the demographies in one grouptype changes patterns of kinship dynamics in both grouptypes, Fig. A4-A5). Such population-scale coupling means that predictions of individuals’ age-specific relatedness in populations with group size local variation cannot be obtained by interpolating those by existing models assuming no group size local variation (e.g., Fig. A3), because the underlying processes by which such coupling is played out differ between group-size homogeneous versus heterogeneous populations (e.g., Fig. A1-A2, A6-A7). In summary, our incorporation of group size local variation does more than extend previous group-size homogeneous models — it shows that, within a genetically connected meta-population, the effects of sex, age, dispersal, and mating on an individual’s average relatedness to its groupmates cannot be understood by examining each local grouptype in isolation whilst coexisting.

The qualitative patterns of kinship dynamics for both sexes across the social mammal cases by our model, and the way they relate to the underlying demographic processes, are the same for both smaller and larger groups. Differences in local kinship dynamics nevertheless result from the asymmetric impacts of these demographic processes between smaller and larger groups. In the presence of group size local variation within a genetically connected population, larger groups tend to preserve more local ancestry: newly recruited breeders (either sex) are relatively more likely to be natively-born, and locally born offspring are more likely to be sired by local males (when larger groups contain more breeders of the relevant sex — see analytical proofs on p. 40 but also the impacts of breeder sex ratio on p. 49 in the appendix); while higher probabilities of local recruiting and siring strengthen local relatedness (by increasing the chance of individuals sharing local ancestry), we show that an individual’s relatedness to others is consistently lower in larger groups across the three typical social mammal systems explored.

This apparent paradox arises because two opposing mechanisms act simultaneously: while lower local recruiting and siring means introducing relatively more unrelated lineages into smaller groups, reducing the probability of identity by descent (i.e., diluting local relatedness), smaller groups also contain fewer breeders, so two individuals are much more likely to trace their ancestry back to the same parents (i.e., stronger genetic drift, as the probability of lineage coalescence scales inversely with the numbers of breeders — see recursions in the appendix). Whether smaller groups show higher relatedness therefore depends on the relative strengths of faster lineage coalescence versus greater ancestry dilution. Our results across the three social mammal contexts thus suggest that, when these opposing mechanisms are combined, the relatively stronger coalescence in smaller groups outweighs local relatedness dilution by relatively higher gene inflow (i.e., lower local recruiting and/or siring probabilities), yielding higher relatedness therein. Interestingly, such outweighing is consistent across these social mammal contexts, despite they are characterized by distinct patterns dispersal and local mating. However, this consistency should not be interpreted as being universal, as the opposite can be true under other demographic regimes (e.g., in systems with moderate local mating and dispersal of the sexes, males in smaller groups are consistently less related to their groups, Fig. A4; see also Fig. A5).

Our study also provide new insights into the theory of social life history evolution, by showing how locally varying kinship environments can generate and explain variations in age-linked trends in social behaviors, such as within-population variation in the timing of shifts from harming to helping (i.e., social life history traits such as menopause in social mammals, Nielsen et al., 2021; Thomas et al., 2001). However, here we also highlight the empirical challenges of testing these evolutionary predictions in natural populations. Particularly, a common yet key complication that comes into play is group size variation itself: natural group sizes often fluctuate across ecological and evolutionary timescales, making it empirically difficult to identify consistently smaller from larger groups (especially in fission-fusion societies where group size and membership change frequently and dramatically), whereas with the scope focused on the qualitative, expected effects of smaller versus larger groups on local kinship dynamics and selection gradients for helping/harming, our model assumes persistent, deterministic variation in local group size (i.e., group size is treated as a local demographic state variable instead of a random variable). While the expected effects of group size on kinship dynamics may be detectable in clearly demographically and socially structured populations, inferring the deeper evolutionary causal link between group size and age-related trends in social traits is more challenging.

We highlight two future directions to address this. First, theoretical work could allow group size to fluctuate (through births, deaths, dispersal, fission and fusion, etc.) to make it possible to ask how group size temporal variation, rather than persistent structural variation alone, shapes selection on behaviors. In addition, incorporating age-specific variation in survival and reproduction could link group size more explicitly to kin-selected behavioral trends across the full biological lifespan. Second, empirically linking group size to age-linked trends in behavioral traits requires establishing and/or maintaining long-term study systems for individual-level longitudinal data on demography, kinship, and behaviors. With existing datasets, however, a key step is to identify the social scale at which group size variation most reliably maps onto variations in kinship dynamics and age-linked behavioral trends (e.g., broader analyses may contrast smaller and larger subpopulations while finer-scale analyses may contrast smaller and larger groups within a single subpopulation instead). To illustrate the empirical challenges and uncertainties associated with this social scaling, we draw on facts revealed by long-term census datasets from three killer whale populations.

At the broadest social scale the data permits, comparisons among the northern resident, the southern resident, and the transient populations [24] shows that 1) resident populations typically form larger pods than the transients, and 2) transient females achieve 95% of their lifetime reproductive outputs earlier than females in the resident populations (Nielsen et al., 2021). This pattern is qualitatively consistent with our prediction that smaller social groups should be associated with earlier selective shifts from harming to helping; however, inference at this scale is constrained not only by the very small sample size (i.e., 3 populations), which precludes strong statistical test, but also by the clear ecological differences between the resident and transient ecotypes, which confound any causal interpretation of group size effects. Thus, beyond small sample size, this example demonstrates the challenge of disentangling group size from correlated ecological and evolutionary factors. At a finer social scale, matrilines form the core social units within each population, while individuals from different matrilines associate into stable social clusters as pods (Bigg et al., 1990; Ellis et al., 2022). Using long-term, individually-explicit census data available from the Southern resident population comprising three stable pods (Wiles, 2016), we examined associations between pod size, age class, and female reproductive strategy (see appendix for details). Estimated age-class- and pod-specific differences in annual birth probability were small, with slightly higher predicted birth probabilities for younger than older mothers, particularly in smaller pods (Fig. A8); while this pattern is again qualitatively consistent with our prediction (smaller groups favor earlier shifts from female harming to helping), inference at this scale remains constrained by only three pods and restricted variation in pod size in a single population. Consequently, substantial uncertainty remains on how much these observations reflect underlying effects of age and pod size on reproductive strategies (though the dataset provides a foundation for future rigorous tests of our predictions).

In conclusion, while empirically testing the predicted patterns of age-linked trends in social behaviors in relation to group size remains challenging, we suggest that group size local variation is a fundamental component underlying the diverse kinship environments individuals of both sexes experience as they age in natural populations, and such variations may therefore help to explain widespread variations in social traits, including the evolution and timing of menopause in social mammals (Nielsen et al., 2021; Thomas et al., 2001).

## DATA AVAILABILITY

The expressions calculating sex-age-specific relatedness and selection gradients are given in the appendix. All the code and data used are available at https://github.com/ecopeng/GroupSize_KinshipDynamics_Selection.

## AUTHORS’ CONTRIBUTIONS

PH and RAJ led the conceptualization, modelling, and writing for this study with supports and inputs from other co-authors. MNW, SE and DPC led the data curation, analysis, and interpretation. DWF, MAC, DPC and RAJ contributed to acquiring funding for this study. All authors gave final approval for the publication of this study.

## ACKNOWLEDGEMENTS

We thank Andy Gardner and four anonymous reviewers for their constructive comments, and the Natural Environmental Research Council for granting us the funding (No. NE/S010327/1) to conduct this study.

