## APPENDIX for "Group size modulates kinship dynamics and selection on social traits"

**1 THE BASELINE MODEL OF KINSHIP DYNAMICS** **4**

**5 COMPARISON WITH HOMOGENEOUS POPULATIONS** **44**

**6 GROUP-SIZE-SPECIFIC RELATEDNESS CANNOT BE INTERPOLATED** **47**

**7 EXPLORING DEMOGRAPHIC EFFECTS ON KINSHIP DYNAMICS UNDER GROUP SIZE LOCAL VARIATION** **49**

**9 FEMALE KILLER WHALE REPRODUCTIVE RATE** **59**

#### TABLE OF NOTATIONS

| symbol | meaning |
|---|---|
| $\alpha, \beta$ | grouptypes ($\alpha$: smaller group, $\beta$: larger group; group size is total breeder number) |
| $f, m$ | sex labels ($f$: female, $m$: male) |
| $u$ | proportion of $\alpha$-type (smaller) groups (larger or $\beta$-type groups: $1-u$) |
| $n_f, n_m$ | numbers of female and male breeders per group in the baseline model (group-size homogeneous) |
| $n_{f\alpha}, n_{m\alpha}$ | numbers of female and male breeders in an $\alpha$-type group |
| $n_{f\beta}, n_{m\beta}$ | numbers of female and male breeders in a $\beta$-type group |
| $\bar{n}_f$ | frequency-weighted average number of female breeders per group: $\bar{n}_f = un_{f\alpha} + (1-u)n_{f\beta}$ |
| $\bar{n}_m$ | frequency-weighted average number of male breeders per group: $\bar{n}_m = un_{m\alpha} + (1-u)n_{m\beta}$ |
| $\bar{n}$ | frequency-weighted average group size: $\bar{n} = u(n_{f\alpha} + n_{m\alpha}) + (1-u)(n_{f\beta} + n_{m\beta})$ |
| $p$ | offspring production per female breeder per timestep |
| $x$ | primary sex ratio (fraction of female offspring) |
| $d_f, d_m$ | female and male offspring dispersal probabilities |
| $\mu_f, \mu_m$ | female and male breeder mortality probabilities per timestep |
| $m$ | baseline local siring probability for an offspring born in a group |
| $\pi_{f\alpha}, \pi_{m\alpha}$ | probabilities that a female/male candidate or newly recruited breeder in an $\alpha$-type group is native |
| $\pi_{f\beta}, \pi_{m\beta}$ | probabilities that a female/male candidate or newly recruited breeder in a $\beta$-type group is native |
| $m_\alpha, m_\beta$ | probabilities that an offspring born in an $\alpha$- or $\beta$-type group is sired by a local male |
| $a$ | breeder age (since recruitment into the breeding pool — not absolute biological age) |
| $g_f, g_m$ | PIBDs for two homologous alleles sampled from the same female/male breeder |
| $g_{ff}, g_{mm}$ | PIBDs for two homologous alleles sampled from two distinct female/male breeders in a group |
| $g_{fm} = g_{mf}$ | PIBD for two homologous alleles sampled from a female and a male breeder in a group |
| $g'_f, g'_m, g'_{ff}, g'_{fm}, g'_{mf}, g'_{mm}$ | corresponding PIBDs after one timestep |
| $g^0_{ff}, g^0_{fm}, g^0_{mf}, g^0_{mm}$ | PIBDs for a newly recruited focal breeder and a distinct groupmate in given sex ($1^{st}/2^{nd}$ subscript: groupmate/focal sex) |
| $g^a_{ff}, g^a_{fm}, g^a_{mf}, g^a_{mm}$ | PIBDs for a focal breeder aged $a$ ($\geq 1$) and a groupmate ($1^{st}/2^{nd}$ subscript: groupmate/focal sex) |
| $g^{\alpha a}_{ff}, g^{\alpha a}_{fm}, g^{\alpha a}_{mf}, g^{\alpha a}_{mm}$ | PIBDs for a focal breeder aged $a$ and a groupmate in an $\alpha$-type group ($1^{st}/2^{nd}$ subscript: groupmate/focal sex) |
| $g^{\beta a}_{ff}, g^{\beta a}_{fm}, g^{\beta a}_{mf}, g^{\beta a}_{mm}$ | PIBDs for a focal breeder aged $a$ and a groupmate in a $\beta$-type group ($1^{st}/2^{nd}$ subscript: groupmate/focal sex) |
| $g^{\alpha a}_f, g^{\beta a}_f$ | weighted average PIBD of a focal female aged $a$ to all groupmates in an $\alpha$- or $\beta$-type group |
| $g^{\alpha a}_m, g^{\beta a}_m$ | weighted average PIBD of a focal male aged $a$ to its groupmates in an $\alpha$- or $\beta$-type group |
| $r^{\alpha a}_{ff}, r^{\alpha a}_{fm}, r^{\alpha a}_{mf}, r^{\alpha a}_{mm}$ | relatedness coef (normalized PIBDs): a breeder aged $a$ and others in an $\alpha$-group ($1^{st}/2^{nd}$ subscript: groupmate/focal sex) |
| $r^{\beta a}_{ff}, r^{\beta a}_{fm}, r^{\beta a}_{mf}, r^{\beta a}_{mm}$ | relatedness coef (normalized PIBDs): a breeder aged $a$ and others in a $\beta$-group ($1^{st}/2^{nd}$ subscript: groupmate/focal sex) |
| $\boldsymbol{W}$ | matrix of expected (neutral, autosomal) allele copy transmission among sex-grouptype classes |
| $w_{i\leftarrow j}$ | entry of $\boldsymbol{W}$: expected number of descendant allele copies in destination class $i$ from a copy in source class $j$ |
| $\boldsymbol{z}_t$ | vector of expected allele copy abundances in sex-grouptype classes at timestep $t$ |
| $z^t_{f\alpha}, z^t_{m\alpha}, z^t_{f\beta}, z^t_{m\beta}$ | entries of $\boldsymbol{z}_t$ (expected allele copy abundance in each sex-grouptype class) |
| $\lambda$ | dominant eigenvalue of $\boldsymbol{W}$ ($\lambda = 1$ under allele copy transmission stationarity) |
| $\boldsymbol{v}$ | dominant (left) eigenvector satisfying $\boldsymbol{vW} = \boldsymbol{v}$ to derive sex-grouptype-specific reproductive values |
| $v_{f\alpha}, v_{m\alpha}, v_{f\beta}, v_{m\beta}$ | unscaled sex-grouptype-specific reproductive values |
| $\bar{v}$ | average reproductive value weighted by relative breeder abundance across sex-grouptype classes |
| $v'_{f\alpha}, v'_{m\alpha}, v'_{f\beta}, v'_{m\beta}$ | scaled reproductive values across sex-grouptype classes |
| $w_{f\alpha}, w_{m\alpha}, w_{f\beta}, w_{m\beta}$ | fitness functions for an allele borne by a focal breeder in each sex-grouptype class (subscripts without $\leftarrow$) |
| $p_{if\alpha}, p_{if\beta}$ | fecundity of a focal female breeder in an $\alpha$- or $\beta$-type group |
| $p_{im\alpha}, p_{im\beta}$ | reproductive output (or mating success) of a focal male breeder in an $\alpha$- or $\beta$-type group |
| $p_{gf\alpha}, p_{gf\beta}$ | average fecundity of female breeders in the focal group of given type |
| $p_{gm\alpha}, p_{gm\beta}$ | average reproductive output (or mating success) of male breeders in the focal group of given type |
| $p^*_{gf\alpha}, p^*_{gf\beta}$ | excluding-focal average fecundity of female breeders in the focal group of given type |
| $p^*_{gm\alpha}, p^*_{gm\beta}$ | excluding-focal average reproductive output (or mating success) of male breeders in the focal group of given type |
| $p_f, p_m$ | population-level average female fecundity and male reproductive output (or mating success) |

| symbol | meaning |
|---|---|
| $\mu_{if\alpha}, \mu_{if\beta}$ | mortality of a focal female breeder in an $\alpha$- or $\beta$-type group |
| $\mu_{im\alpha}, \mu_{im\beta}$ | mortality of a focal male breeder in an $\alpha$- or $\beta$-type group |
| $\mu_{gf\alpha}, \mu_{gf\beta}$ | average mortality of female breeders in the focal grouptype |
| $\mu_{gm\alpha}, \mu_{gm\beta}$ | average mortality of male breeders in the focal grouptype |
| $\mu^*_{gf\alpha}, \mu^*_{gf\beta}$ | excluding-focal group average mortality of female breeders in the focal grouptype |
| $\mu^*_{gm\alpha}, \mu^*_{gm\beta}$ | excluding-focal group average mortality of male breeders in the focal grouptype |
| $b$ | total marginal (fecundity/mortality) benefit/cost imposed on recipients (sign determines helping vs harming) |
| $c$ | actor's marginal personal cost (fecundity/mortality) |
| $b^*$ | critical value of $b$ at which the net inclusive-fitness effect of the behaviour is zero |
| $c/b^*$ | critical ratio measuring direction and strength of selection; positive/negative favors helping/harming |
| $F^{\alpha a}_{ff}, F^{\alpha a}_{mf}, F^{\alpha a}_{fm}, F^{\alpha a}_{mm}$ | inclusive fitness effects of behaviours with fecundity consequences in an $\alpha$-group ($1^{st}/2^{nd}$ subscript: recipient/focal sex) |
| $F^{\beta a}_{ff}, F^{\beta a}_{mf}, F^{\beta a}_{fm}, F^{\beta a}_{mm}$ | inclusive fitness effects of behaviours with fecundity consequences in a $\beta$-group ($1^{st}/2^{nd}$ subscript: recipient/focal sex) |
| $M^{\alpha a}_{ff}, M^{\alpha a}_{mf}, M^{\alpha a}_{fm}, M^{\alpha a}_{mm}$ | inclusive fitness effects of behaviours with mortality consequences in an $\alpha$-group ($1^{st}/2^{nd}$ subscript: recipient/focal sex) |
| $M^{\beta a}_{ff}, M^{\beta a}_{mf}, M^{\beta a}_{fm}, M^{\beta a}_{mm}$ | inclusive fitness effects of behaviours with mortality consequences in a $\beta$-group ($1^{st}/2^{nd}$ subscript: recipient/focal sex) |
| $F^{\alpha a}_{f}, F^{\beta a}_{f}$ | weighted average fecundity consequences for a random groupmate of a focal female aged $a$ in an $\alpha$- or $\beta$-type group |
| $M^{\alpha a}_{m}, M^{\beta a}_{m}$ | weighted average mortality consequences for a random groupmate of a focal male aged $a$ in an $\alpha$- or $\beta$-type group |

### 1 THE BASELINE MODEL OF KINSHIP DYNAMICS

In the model, the sex- and age-specific average relatedness of individuals to others of given sex in smaller and larger social groups in a subdivided, genetically connected population is calculated by developing the baseline analytical model of kinship dynamics briefly described as below (see more details of the model in Johnstone and Cant, 2008, 2010, Ellis et al., 2022).

#### 1.1 Demographic Assumptions

In the *baseline* model (e.g., Ellis et al., 2022), individuals (i.e., established breeders who occupy limited breeding vacancies in groups) are assumed diploid, sexual, and inhabit in an infinite number of groups (a.k.a. 'islands', demes or patches, connected by gene flows), each of which is with $n_f$ female and $n_m$ male breeders (i.e., in this model total patch or group size is the same for all). At each discrete timestep (i.e., from a given age to the next), each female breeder (i.e., age $\geq 1$) in each group reproduces a sufficiently large number, say $p$, of offspring consisting a given fraction, say $x$, of females while the rest $1 - x$ are males. The probability that an offspring born in a group is sired by a male breeder from the same group is $m$ (i.e., probability of being derived from local mating, a measure of the mating success of local males), while the probability of being sired by a male breeder from elsewhere is $1 - m$. After births, each female offspring disperses to a random (non-natal) group with probability $d_f$, so does each male offspring in each group with the probability $d_m$. The female and male offspring that did not disperse in each group, together with the pool of immigrants from the population, compete in a 'fair lottery' manner for the breeding vacancies (of their own sexes) created by the deaths of those female and male breeders in each group — at each timestep each female breeder dies with the probability $\mu_f$, as each male breeder does so with the probability $\mu_m$. Offspring that failed to obtain breeding vacancies die, and such demographic processes (i.e., mating, birth, death, dispersal, competition) repeat.

#### 1.2 Local Relatedness

With the above assumptions, the expected genetic relatedness between individuals in a group, defined as the probability of identity by descent (PIBD) of two homologous neutral alleles (or genes) — each of which is sampled from each of the individuals, can be derived. Let $g_f$ and $g_m$ denote the PIBDs of two such alleles repeatedly sampled from the same female and male breeder, respectively. Let $g_{ff}$ or $g_{mm}$ denote the PIBDs of two such alleles, each of which is sampled from a distinct female or male breeder in a group, respectively, while $g_{fm} = g_{mf}$ denote the PIBDs of two such alleles, one of which is sampled from a female breeder while the other from a male breeder. Similarly, let $g_f^{'}$, $g_m^{'}$, $g_{ff}^{'}$, $g_{fm}^{'} = g_{mf}^{'}$ and $g_{mm}^{'}$ denote such PIBDs of two such alleles sampled at the next timestep in a group. Then, these PIBDs can be described as below (see details on the derivations in Johnstone and Cant, 2008, 2010):

$$g_f^{'} = (1-\mu_f)\,g_f + \mu_f\left[m\frac{1}{2}\,(1+g_{fm}) + (1-m)\,\frac{1}{2}\right] \tag{1}$$

$$g_m^{'} = (1-\mu_m)\,g_m + \mu_m\left[m\frac{1}{2}\,(1+g_{fm}) + (1-m)\,\frac{1}{2}\right] \tag{2}$$

$$\begin{aligned} g_{ff}^{'} = {} & \left(1-\mu_f\right)^2 g_{ff} + \\ & 2\left(1-\mu_f\right)\mu_f\left(1-d_f\right)\left\{m\frac{1}{2}\left[\frac{1}{n_f}g_f + \left(1-\frac{1}{n_f}\right)g_{ff} + g_{fm}\right] + \right. \\ & \left. (1-m)\,\frac{1}{2}\left[\frac{1}{n_f}g_f + \left(1-\frac{1}{n_f}\right)g_{ff}\right]\right\} + \\ & \mu_f^2\left(1-d_f\right)^2\left\{m^2\frac{1}{4}\left[\frac{1}{n_f}g_f + \left(1-\frac{1}{n_f}\right)g_{ff} + 2g_{fm} + \frac{1}{n_m}g_m + \left(1-\frac{1}{n_m}\right)g_{mm}\right] + \right. \\ & 2m\,(1-m)\,\frac{1}{4}\left[\frac{1}{n_f}g_f + \left(1-\frac{1}{n_f}\right)g_{ff} + g_{fm}\right] + \\ & \left. (1-m)^2\,\frac{1}{4}\left[\frac{1}{n_f}g_f + \left(1-\frac{1}{n_f}\right)g_{ff}\right]\right\} \end{aligned} \tag{3}$$

$$
\begin{aligned}
g'_{fm} = g'_{mf} \\
&= \left(1-\mu_f\right)(1-\mu_m)\, g_{fm} + \\
&\left(1-\mu_f\right)\mu_m (1-d_m)\left\{ m\frac{1}{2}\left[\frac{1}{n_f}g_f + \left(1-\frac{1}{n_f}\right)g_{ff} + g_{fm}\right] + \right. \\
&\left. (1-m)\frac{1}{2}\left[\frac{1}{n_f}g_f + \left(1-\frac{1}{n_f}\right)g_{ff}\right]\right\} + \\
&\mu_f (1-\mu_m)\left(1-d_f\right)\left\{ m\frac{1}{2}\left[g_{fm} + \frac{1}{n_m}g_m + \left(1-\frac{1}{n_m}\right)g_{mm}\right] + \right. \\
&\left. (1-m)\frac{1}{2}g_{fm}\right\} + \\
&\mu_f\mu_m\left(1-d_f\right)(1-d_m)\left\{ m^2\frac{1}{4}\left[\frac{1}{n_f}g_f + \left(1-\frac{1}{n_f}\right)g_{ff} + 2g_{fm} + \frac{1}{n_m}g_m + \left(1-\frac{1}{n_m}\right)g_{mm}\right] + \right. \\
&2m(1-m)\frac{1}{4}\left[\frac{1}{n_f}g_f + \left(1-\frac{1}{n_f}\right)g_{ff} + g_{fm}\right] + \\
&\left. (1-m)^2\frac{1}{4}\left[\frac{1}{n_f}g_f + \left(1-\frac{1}{n_f}\right)g_{ff}\right]\right\} \qquad (4)
\end{aligned}
$$

$$
\begin{aligned}
g'_{mm} = (1-\mu_m)^2 g_{mm} + \\
&2(1-\mu_m)\mu_m(1-d_m)\left\{ m\frac{1}{2}\left[g_{fm} + \frac{1}{n_m}g_m + \left(1-\frac{1}{n_m}\right)g_{mm}\right] + \right. \\
&\left. (1-m)\frac{1}{2}g_{fm}\right\} + \\
&\mu_m^2(1-d_m)^2\left\{ m^2\frac{1}{4}\left[\frac{1}{n_f}g_f + \left(1-\frac{1}{n_f}\right)g_{ff} + 2g_{fm} + \frac{1}{n_m}g_m + \left(1-\frac{1}{n_m}\right)g_{mm}\right] + \right. \\
&2m(1-m)\frac{1}{4}\left[\frac{1}{n_f}g_f + \left(1-\frac{1}{n_f}\right)g_{ff} + g_{fm}\right] + \\
&\left. (1-m)^2\frac{1}{4}\left[\frac{1}{n_f}g_f + \left(1-\frac{1}{n_f}\right)g_{ff}\right]\right\} \qquad (5)
\end{aligned}
$$

For example, the derivation of $g_f$ [equation (1)] or $g_m$ [equation (2)] can be illustrated with the flowchart below:

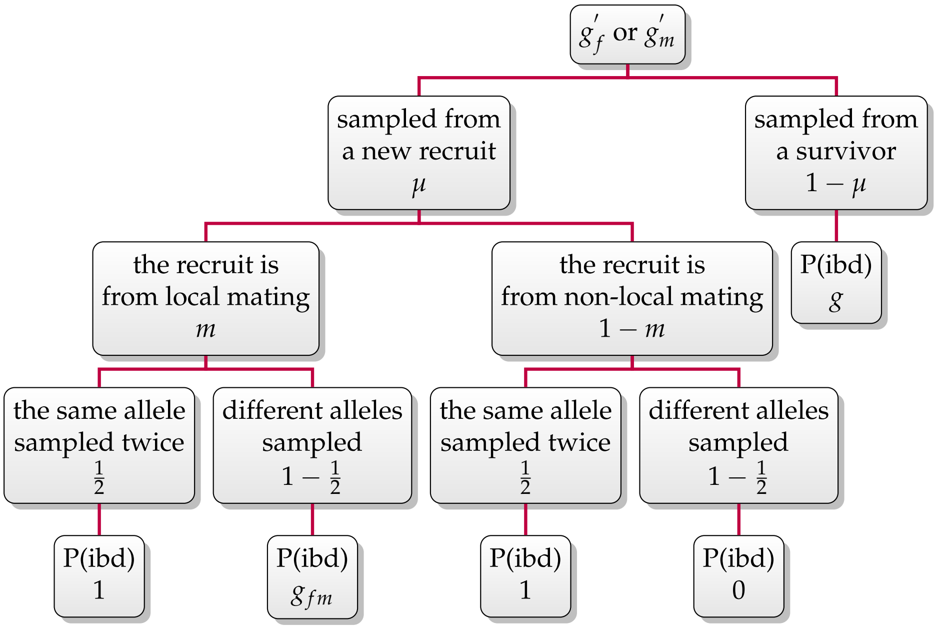

For another example, $g_{ff}$ [equation (3)] can be derived following the flowchart given as

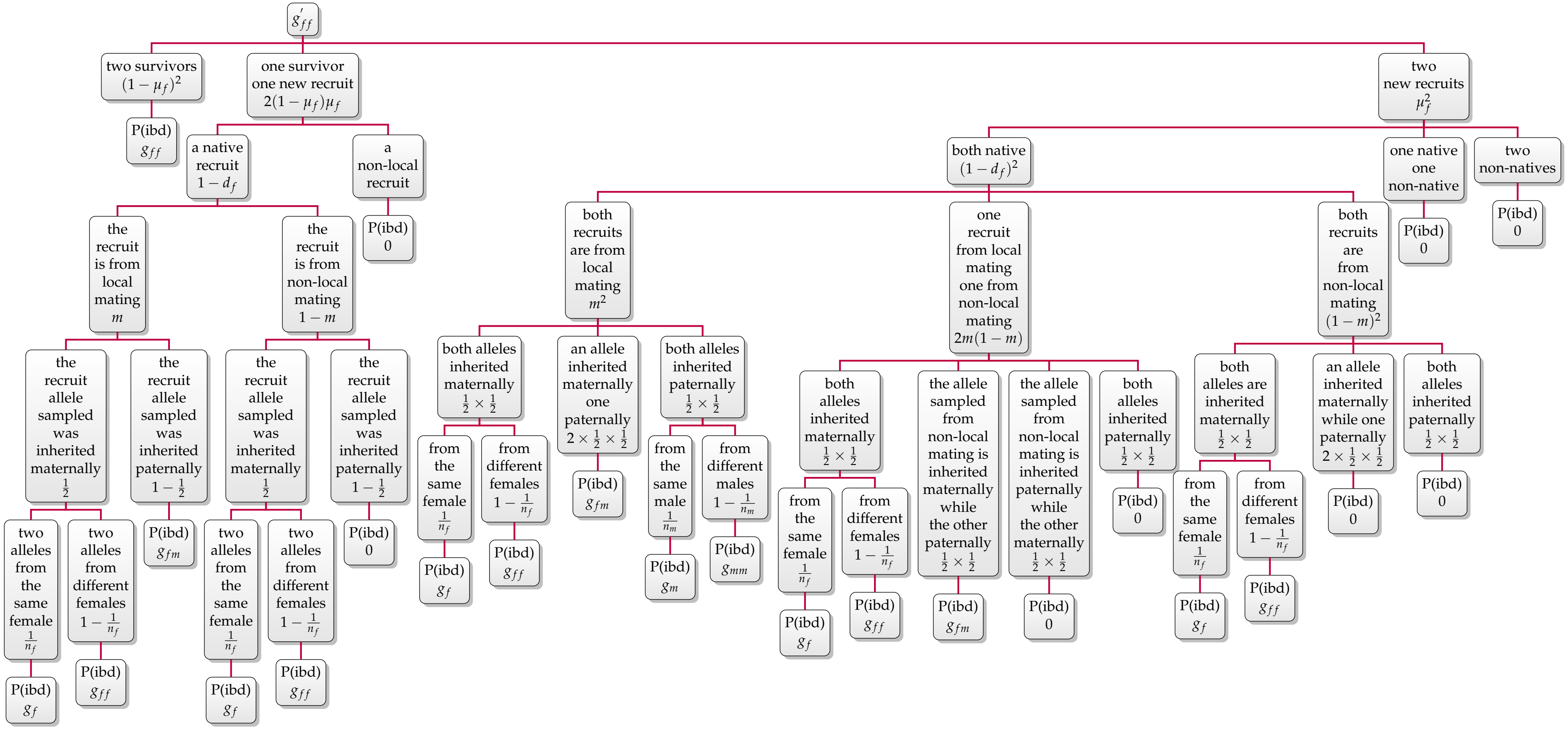

When the PIBDs of two such alleles sampled at a given timestep remain what they were in the last timestep (i.e., when $g' = g$ in each of the above equations), the relatedness between breeders in each group reaches the equilibrium, and by solving such equations, we get these PIBDs as *sex-specific* relatedness in each group.

##### 1.3 Sex- and Age-specific Relatedness

Now we can derive the expected relatedness between a focal breeder with given *sex* and *age* to a randomly sampled groupmate with given *sex* (but implicit age). Let $g^0_{fm}$ denote the PIBD of two such alleles, given that one is sampled from a focal male breeder aged 0 in a group (i.e., a newly recruited male breeder in the group as the focal individual indicated by the last letter '$m$' in the subscripts of $g^0_{fm}$; note that here the age '0' is defined in a relative sense: it refers to the *youngest breeders* who just obtained breeding vacancies in a group, rather than the absolute age of the offspring in a group defined in the main texts), while the other is sampled from a female breeder (age-implicit) in the patch. Similarly, let $g^0_{ff}$, $g^0_{mf}$ and $g^0_{mm}$ denote such PIBDs under other combinations of the sexes of the two breeders sampled (where the youngest breeder in focus is with given sex and aged '0', while the other is with given sex but age-implicit). Then, the expected relatedness of a newly recruited focal breeder to others in a group can be described as:

$$
\begin{aligned}
g^0_{ff} = \left(1-\mu_f\right)\left(1-d_f\right)\Bigg\{ & m\frac{1}{2}\left[\frac{1}{n_f}g_f + \left(1-\frac{1}{n_f}\right)g_{ff} + g_{fm}\right] + \\
& (1-m)\frac{1}{2}\left[\frac{1}{n_f}g_f + \left(1-\frac{1}{n_f}\right)g_{ff}\right]\Bigg\} + \\
\left(1-d_f\right)\mu_f\left(1-d_f\right)\Bigg\{ & m\Bigg[m\frac{1}{4}\left(\frac{1}{n_f}g_f + \left(1-\frac{1}{n_f}\right)g_{ff} + 2g_{fm} + \frac{1}{n_m}g_m + \left(1-\frac{1}{n_m}\right)g_{mm}\right) + \\
& (1-m)\frac{1}{4}\left(\frac{1}{n_f}g_f + \left(1-\frac{1}{n_f}\right)g_{ff} + g_{fm}\right)\Bigg] + \\
& (1-m)\Bigg[m\frac{1}{4}\left(\frac{1}{n_f}g_f + \left(1-\frac{1}{n_f}\right)g_{ff} + g_{fm}\right) + \\
& (1-m)\frac{1}{4}\left(\frac{1}{n_f}g_f + \left(1-\frac{1}{n_f}\right)g_{ff}\right)\Bigg]\Bigg\}
\end{aligned}
\tag{6}
$$

$$
\begin{aligned}
g_{fm}^{0} = \left(1-\mu_f\right)\left(1-d_m\right)\Bigg\{ & m\frac{1}{2}\left[\frac{1}{n_f}g_f+\left(1-\frac{1}{n_f}\right)g_{ff}+g_{fm}\right]+ \\
& (1-m)\frac{1}{2}\left[\frac{1}{n_f}g_f+\left(1-\frac{1}{n_f}\right)g_{ff}\right]\Bigg\}+ \\
\left(1-d_f\right)\mu_f\left(1-d_m\right)\Bigg\{ & m\Bigg[m\frac{1}{4}\left(\frac{1}{n_f}g_f+\left(1-\frac{1}{n_f}\right)g_{ff}+2g_{fm}+\frac{1}{n_m}g_m+\left(1-\frac{1}{n_m}\right)g_{mm}\right)+ \\
& (1-m)\frac{1}{4}\left(\frac{1}{n_f}g_f+\left(1-\frac{1}{n_f}\right)g_{ff}+g_{fm}\right)\Bigg]+ \\
& (1-m)\Bigg[m\frac{1}{4}\left(\frac{1}{n_f}g_f+\left(1-\frac{1}{n_f}\right)g_{ff}+g_{fm}\right)+ \\
& (1-m)\frac{1}{4}\left(\frac{1}{n_f}g_f+\left(1-\frac{1}{n_f}\right)g_{ff}\right)\Bigg]\Bigg\}
\end{aligned} \tag{7}
$$

$$
\begin{aligned}
g_{mf}^{0} = \left(1-\mu_m\right)\left(1-d_f\right)\Bigg\{ & m\frac{1}{2}\left[g_{fm}+\frac{1}{n_m}g_m+\left(1-\frac{1}{n_m}\right)g_{mm}\right]+ \\
& (1-m)\frac{1}{2}g_{fm}\Bigg\}+ \\
\left(1-d_m\right)\mu_m\left(1-d_f\right)\Bigg\{ & m\Bigg[m\frac{1}{4}\left(\frac{1}{n_f}g_f+\left(1-\frac{1}{n_f}\right)g_{ff}+2g_{fm}+\frac{1}{n_m}g_m+\left(1-\frac{1}{n_m}\right)g_{mm}\right)+ \\
& (1-m)\frac{1}{4}\left(\frac{1}{n_f}g_f+\left(1-\frac{1}{n_f}\right)g_{ff}+g_{fm}\right)\Bigg]+ \\
& (1-m)\Bigg[m\frac{1}{4}\left(\frac{1}{n_f}g_f+\left(1-\frac{1}{n_f}\right)g_{ff}+g_{fm}\right)+ \\
& (1-m)\frac{1}{4}\left(\frac{1}{n_f}g_f+\left(1-\frac{1}{n_f}\right)g_{ff}\right)\Bigg]\Bigg\}
\end{aligned} \tag{8}
$$

$$
\begin{aligned}
g^0_{mm} = (1-\mu_m)(1-d_m)\Bigg\{ & m\frac{1}{2}\left[\frac{1}{n_m}g_m + \left(1-\frac{1}{n_m}\right)g_{mm} + g_{fm}\right] + \\
& (1-m)\frac{1}{2}g_{fm}\Bigg\} + \\
(1-d_m)\mu_m(1-d_m)\Bigg\{ & m\left[m\frac{1}{4}\left(\frac{1}{n_f}g_f + \left(1-\frac{1}{n_f}\right)g_{ff} + 2g_{fm} + \frac{1}{n_m}g_m + \left(1-\frac{1}{n_m}\right)g_{mm}\right) + \right. \\
& \left.(1-m)\frac{1}{4}\left(\frac{1}{n_f}g_f + \left(1-\frac{1}{n_f}\right)g_{ff} + g_{fm}\right)\right] + \\
& (1-m)\left[m\frac{1}{4}\left(\frac{1}{n_f}g_f + \left(1-\frac{1}{n_f}\right)g_{ff} + g_{fm}\right) + \right. \\
& \left.(1-m)\frac{1}{4}\left(\frac{1}{n_f}g_f + \left(1-\frac{1}{n_f}\right)g_{ff}\right)\right]\Bigg\}
\end{aligned} \tag{9}
$$

Now let $g^a_{fm}$, $g^a_{ff}$, $g^a_{mf}$ and $g^a_{mm}$ denote such PIBDs of the two alleles, one of which is sampled from a focal breeder of given *sex* and *age* $a \geq 1$ while the other is sampled from a distinct breeder with given *sex* but *implicit age* in a group. Then, the average relatedness of a focal individual of given sex and age $a$ to its groupmates can be described by

$$
g^a_{ff} = \left(1-\mu_f\right)g^{a-1}_{ff} + \mu_f\left(1-d_f\right)\left[m\frac{1}{2}g^{a-1}_{mf} + \frac{1}{2}\left(\frac{1}{n_f}g_f + \left(1-\frac{1}{n_f}\right)g^{a-1}_{ff}\right)\right] \tag{10}
$$

$$
g^a_{fm} = \left(1-\mu_f\right)g^{a-1}_{fm} + \mu_f\left(1-d_f\right)\left[m\frac{1}{2}\left(\frac{1}{n_m}g_m + \left(1-\frac{1}{n_m}\right)g^{a-1}_{mm}\right) + \frac{1}{2}g^{a-1}_{fm}\right] \tag{11}
$$

$$
g^a_{mf} = (1-\mu_m)g^{a-1}_{mf} + \mu_m(1-d_m)\left[m\frac{1}{2}g^{a-1}_{mf} + \frac{1}{2}\left(\frac{1}{n_f}g_f + \left(1-\frac{1}{n_f}\right)g^{a-1}_{ff}\right)\right] \tag{12}
$$

$$
g^a_{mm} = (1-\mu_m)g^{a-1}_{mm} + \mu_m(1-d_m)\left[m\frac{1}{2}\left(\frac{1}{n_m}g_m + \left(1-\frac{1}{n_m}\right)g^{a-1}_{mm}\right) + \frac{1}{2}g^{a-1}_{fm}\right] \tag{13}
$$

For example, the derivation of $g^0_{mf}$ [equation (8)] can be illustrated as:

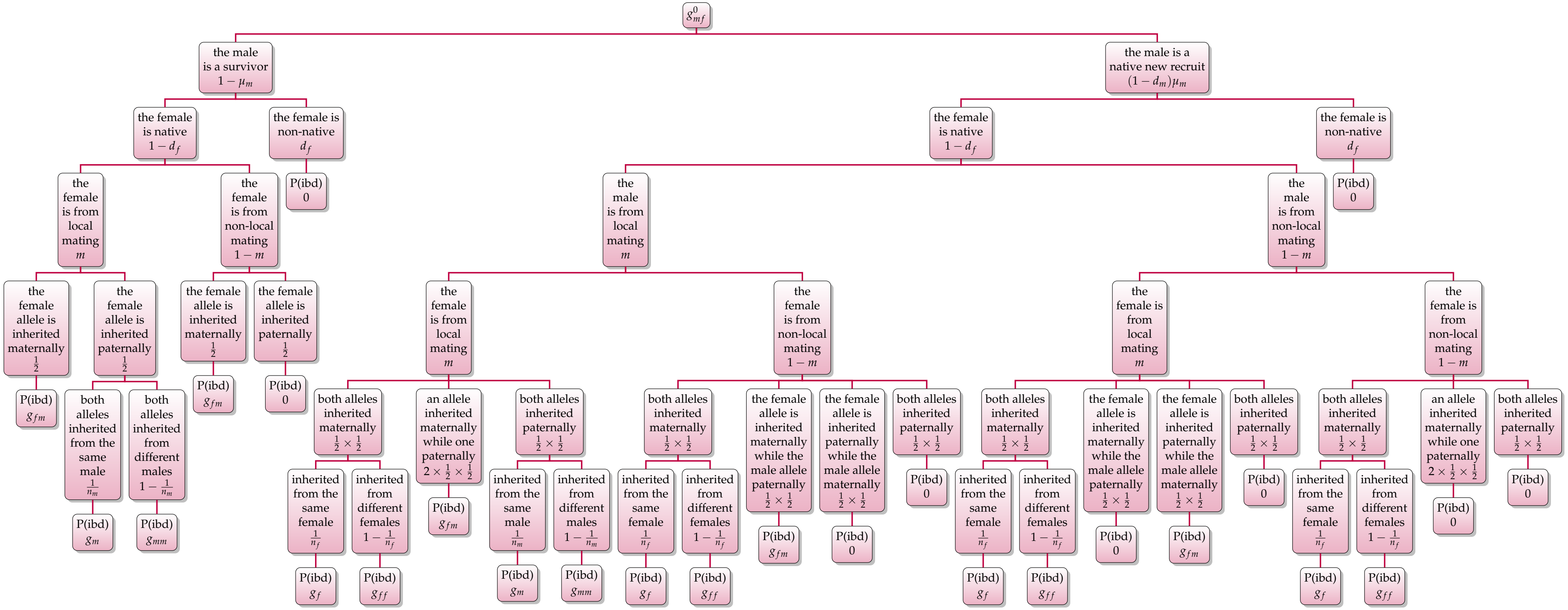

For another example, the derivation of $g^a_{mf}$ [equation (12)] can be illustrated as:

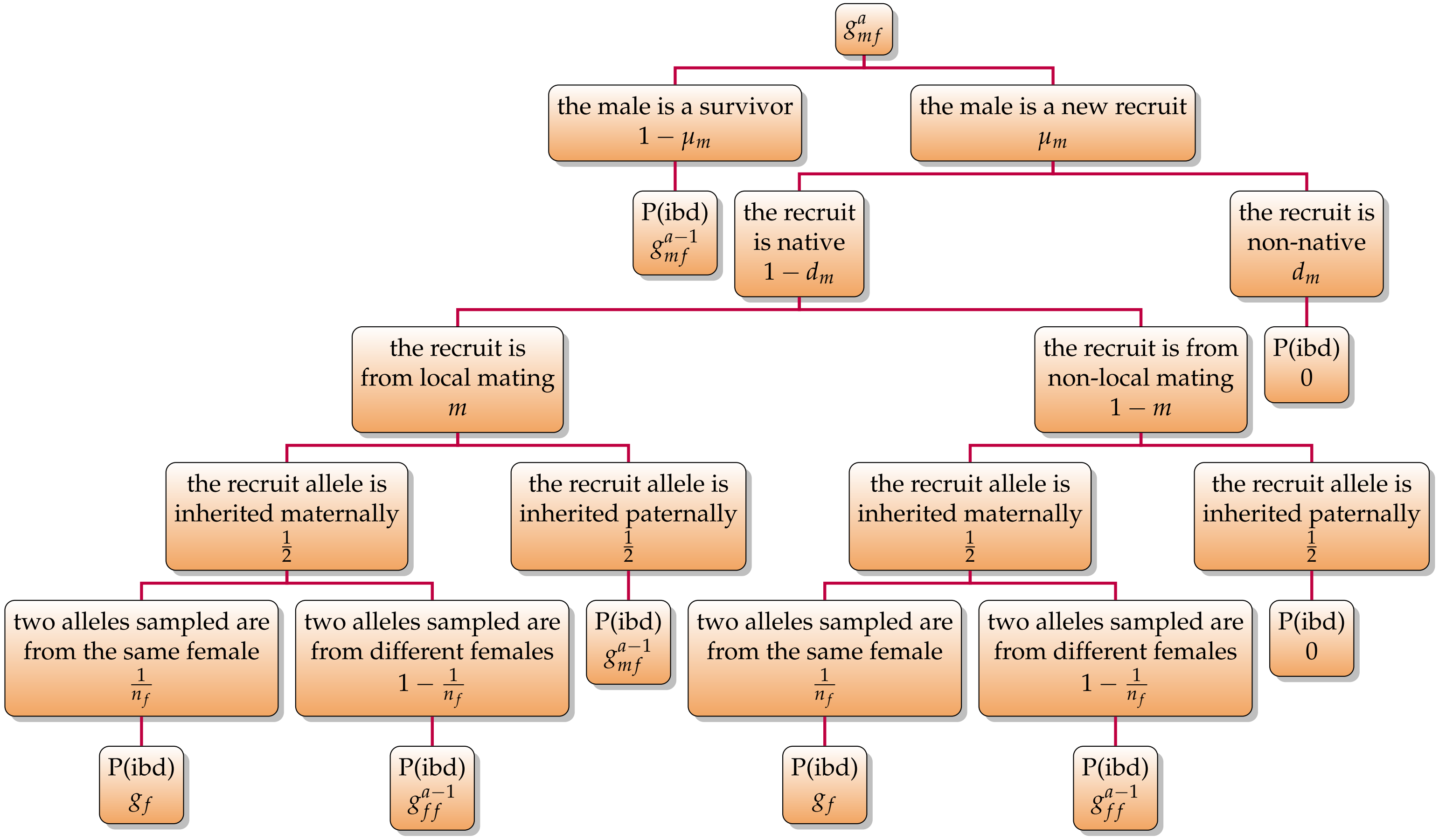

#### 2 EFFECTS OF GROUP SIZE ON KINSHIP DYNAMICS

At this point, with the baseline model of kinship dynamics, we can evaluate the effects of local group size on the sex- and age-specific relatedness of a focal breeder to its groupmates (of given sex) within such a subdivided, yet genetically connected population. Here, we further assume that, following the subdivision of a population, there is a proportion $u$ of the groups are of $\alpha$-type consisting of $n_{f\alpha}$ female and $n_{m\alpha}$ male breeders, while the rest $1-u$ are $\beta$-type groups consisting of $n_{f\beta}$ female and $n_{m\beta}$ male breeders (as in the baseline model, we assume at each timestep each female breeder reproduces a sufficiently large number $p$ of offspring and a fraction $x$ of them are females while the rest are males). As we are interested in group size local variation following population asymmetric division, we assume that $n_{f\alpha}+n_{m\alpha}<n_{f\beta}+n_{m\beta}$ — i.e., the $\alpha$-type groups are relatively smaller than those $\beta$-type groups in terms of their *total size* (otherwise when $n_{f\alpha}=n_{f\beta}$ while $n_{m\alpha}=n_{m\beta}$ our model reduces to the baseline model, and this is how we can generate predictions for a group-size homogeneous population — see details in the `python` script supplemented). Under these assumptions, the average numbers of female and male breeders in a (representative) group in the population are

$$\bar{n}_f = un_{f\alpha}+(1-u)\,n_{f\beta} \tag{14}$$

and

$$\bar{n}_m = un_{m\alpha}+(1-u)\,n_{m\beta} \tag{15}$$

respectively.

##### 2.1 Local Recruiting Probabilities

Then, in an $\alpha$-type group, the probability that a female candidate competing for a local breeding vacancy is native to that group is

$$\pi_{f\alpha}=\frac{(1-d_f)\,xpn_{f\alpha}}{(1-d_f)\,xpn_{f\alpha}+d_f xp\bar{n}_f}$$

$$= \frac{(1-d_f)\, n_{f\alpha}}{(1-d_f)\, n_{f\alpha} + d_f \bar{n}_f} \tag{16}$$

and the corresponding probability for a male candidate is

$$\begin{aligned} \pi_{m\alpha} &= \frac{(1-d_m)\,(1-x)\, p n_{f\alpha}}{(1-d_m)\,(1-x)\, p n_{f\alpha} + d_m\,(1-x)\, p \bar{n}_f} \\ &= \frac{(1-d_m)\, n_{f\alpha}}{(1-d_m)\, n_{f\alpha} + d_m \bar{n}_f} \end{aligned} \tag{17}$$

Because recruitment follows a 'fair lottery' among candidates of the same sex, and the 'lottery success' is independent of whether a candidate is native or immigrant, these same probabilities also give the probability that a newly recruited breeder of the corresponding sex is native to the $\alpha$-type group.

Similarly, in a $\beta$-type group, the probability that a female candidate competing for a local breeding vacancy is native to that group is

$$\begin{aligned} \pi_{f\beta} &= \frac{(1-d_f)\, x p n_{f\beta}}{(1-d_f)\, x p n_{f\beta} + d_f x p \bar{n}_f} \\ &= \frac{(1-d_f)\, n_{f\beta}}{(1-d_f)\, n_{f\beta} + d_f \bar{n}_f} \end{aligned} \tag{18}$$

and the corresponding probability for a male candidate is

$$\begin{aligned} \pi_{m\beta} &= \frac{(1-d_m)\,(1-x)\, p n_{f\beta}}{(1-d_m)\,(1-x)\, p n_{f\beta} + d_m\,(1-x)\, p \bar{n}_f} \\ &= \frac{(1-d_m)\, n_{f\beta}}{(1-d_m)\, n_{f\beta} + d_m \bar{n}_f} \end{aligned} \tag{19}$$

As recruitment follows the same 'fair lottery' process, these probabilities also give the probability that a newly recruited breeder of the corresponding sex is native to the $\beta$-type group.

#### 2.2 Local Siring Probabilities

Meanwhile, the probability that an offspring born in an $\alpha$-type group is sired by a male from that same group is

$$m_{\alpha} = \frac{mn_{m\alpha}}{mn_{m\alpha} + (1-m)\,\bar{n}_m} \tag{20}$$

and the corresponding probability for an offspring born in a $\beta$-type group is

$$m_{\beta} = \frac{mn_{m\beta}}{mn_{m\beta} + (1-m)\,\bar{n}_m} \tag{21}$$

These probabilities also apply to native recruits (i.e., offspring born in a group and later recruited as breeders in that same local group), because siring occurs before dispersal and recruitment, and the 'fair-lottery' recruitment process is independent of paternal origin.

#### 2.3 Expected Sex-, Age-, and Grouptype-specific Relatedness

Then, in a local group of a given *type* (i.e., smaller or larger), the expected relatedness of an individual in a given *sex* and *age* to a random groupmate in any given *sex* can be derived — symbolically we denote, for an example, the relatedness of a focal *female* aged $a$ to a randomly-chosen *male* groupmate in the $\alpha$-type group as $g_{mf}^{\alpha a}$ (the first/second subscript indicates the groupmate/focal sex). Such derivations are achieved by first applying the equations (14)–(21) to those derived in (1)–(5) from the baseline model of kinship dynamics, and solve these equations for the (age-implicit) equilibrium relatedness between individuals (i.e., analogously to setting $g'$ = $g$ in the baseline model of kinship dynamics that does not consider group size local varition), and then applying this same set of equations to those in (6)–(13), to derive an individual's *sex-age-specific* relatedness to a random groupmate — in a target *sex* and *grouptype* (which is used for evaluating the inclussive fitness outcomes of helping/harming; see 3.4). Mathematically, the applications are done by substituting: (I) $1 - d_f$ with either $\pi_{f\alpha}$ or $\pi_{f\beta}$ derived in (16) and (18), respectively, (II) $1 - d_m$ with either $\pi_{m\alpha}$ or $\pi_{m\beta}$ derived in (17) and (19), respectively, and (III) $m$ with either $m_{\alpha}$ or $m_{\beta}$ derived in (20) and (21), respectively.

Lastly, an individual's sex-age-specific average relatedness to its groupmates (both females and males) in a given grouptype can be derived (e.g., Fig. 1 in the main texts). Specifically, the (weighted average) relatedness of a focal *female* aged $a$ to her groupmates in an $\alpha$-type or $\beta$-type group is

$$g^{\alpha a}_{f} = \frac{(n_{f\alpha} - 1) \times g^{\alpha a}_{ff} + n_{m\alpha} \times g^{\alpha a}_{mf}}{n_{f\alpha} - 1 + n_{m\alpha}} \tag{22.1}$$

or

$$g^{\beta a}_{f} = \frac{(n_{f\beta} - 1) \times g^{\beta a}_{ff} + n_{m\beta} \times g^{\beta a}_{mf}}{n_{f\beta} - 1 + n_{m\beta}} \tag{22.2}$$

respectively, while the average relatedness of a focal *male* aged $a$ to his groupmates in an $\alpha$-type or $\beta$-type group is

$$g^{\alpha a}_{m} = \frac{(n_{m\alpha} - 1) \times g^{\alpha a}_{mm} + n_{f\alpha} \times g^{\alpha a}_{fm}}{n_{m\alpha} - 1 + n_{f\alpha}} \tag{22.3}$$

or

$$g^{\beta a}_{m} = \frac{(n_{m\beta} - 1) \times g^{\beta a}_{mm} + n_{f\beta} \times g^{\beta a}_{fm}}{n_{m\beta} - 1 + n_{f\beta}} \tag{22.4}$$

respectively.

#### 3 EFFECTS OF GROUP SIZE ON SELECTION FOR HELPING/HARMING

With the developed model of kinship dynamics taking group size local variation within a population into consideration, we can then examine the extent to which a helping or harming behaviour — with either fecundity or mortality consequences for its bearer and others as recipients, can be favoured by natural selection across the lifespan of individuals in a smaller or larger group following asymmetric subdivision of a population. To achieve this, we first calculate the coefficients of relatedness, and then derive the reproductive values of breeders in given sex and grouptype (smaller or larger), derive the fitness expressions of a mutant allele underlying the social behaviour of the breeders in given sex and grouptype, and lastly, evaluate the inclusive fitness outcomes of the behaviour should it has either fecundity or mortality consequences.

##### 3.1 Coefficient of Relatedness

With an inclusive fitness perspective, the coefficient of relatedness between a focal breeder with given sex and age $a \geq 1$ to a groupmate with given sex can be defined as

$$r_{ff}^{\alpha a} = \frac{g_{ff}^{\alpha a}}{g_f} \tag{23.1}$$

$$r_{fm}^{\alpha a} = \frac{g_{fm}^{\alpha a}}{g_m} \tag{23.2}$$

$$r_{mf}^{\alpha a} = \frac{g_{mf}^{\alpha a}}{g_f} \tag{23.3}$$

$$r_{mm}^{\alpha a} = \frac{g_{mm}^{\alpha a}}{g_m} \tag{23.4}$$

in an $\alpha$-type group, and

$$r_{ff}^{\beta a} = \frac{g_{ff}^{\beta a}}{g_f} \tag{23.5}$$

$$r_{fm}^{\beta a} = \frac{g_{fm}^{\beta a}}{g_m} \tag{23.6}$$

$$r_{mf}^{\beta a} = \frac{g_{mf}^{\beta a}}{g_f} \tag{23.7}$$

$$r_{mm}^{\beta a} = \frac{g_{mm}^{\beta a}}{g_m} \tag{23.8}$$

in a $\beta$-type group. With such a definition, these coefficients of relatedness capture the extent to which the alleles transmitted by a social partner of given sex are related to those transmitted by the focal breeder of given sex and age in a group of given type (i.e., standardized PIBDs as a measure of the 'fidelity' of transmission, Frank, 1998, Rousset, 2004). As before, we use subscripts '$\alpha$' or '$\beta$' to indicate grouptype while '$f$' or '$m$' to indicate sex, such that, for example, $r_{fm}^{\beta a}$ would denote the coefficient of relatedness between a female (with random age) and a focal male aged $a$ in a $\beta$-type (or larger) group (i.e., the $1^{\text{st}}/2^{\text{nd}}$ subscript indicates the groupmate/focal sex).

##### 3.2 Sex-grouptype-specific Reproductive Values

Reproductive values are the expected relative genetic contributions of breeders to a population in the far future (Taylor, 1990, Otto and Day, 2011, Grafen, 2006, Rodrigues and Gardner, 2021). Here, we define reproductive value as the expected relative contribution of a copy of the hypothetical (autosomal, neutral) allele, currently carried by a breeder in each of the four *sex-grouptype classes* (e.g., a female in an $\alpha$-group, a male in a $\beta$-group, etc.), to its future descendant copies in the infinite island population. Briefly, these class-specific reproductive values are a function of (I) the numbers of female and male breeders in a group of given type, (II) the sex-specific dispersal rates of offspring, (III) the rate of local mating, (IV) the prevalence of either grouptype, and (V) the sex-specific mortality (see equations 25.1 - 25.16).

To derive reproductive values, we construct a $4 \times 4$ matrix $\boldsymbol{W}$ to capture the transmissions of the copies of the allele among the classes over one timestep, where each entry $w_{i \leftarrow j}$ gives the *expected* number of the allele copies in the *destination* class $i$ at timestep $t+1$, which are descendant from a copy in the *source* class $j$ at $t$. Specifically, we define $\boldsymbol{W}$ as

$$\boldsymbol{W} = \begin{bmatrix} w_{f\alpha\leftarrow f\alpha} & w_{f\alpha\leftarrow m\alpha} & w_{f\alpha\leftarrow f\beta} & w_{f\alpha\leftarrow m\beta} \\ w_{m\alpha\leftarrow f\alpha} & w_{m\alpha\leftarrow m\alpha} & w_{m\alpha\leftarrow f\beta} & w_{m\alpha\leftarrow m\beta} \\ w_{f\beta\leftarrow f\alpha} & w_{f\beta\leftarrow m\alpha} & w_{f\beta\leftarrow f\beta} & w_{f\beta\leftarrow m\beta} \\ w_{m\beta\leftarrow f\alpha} & w_{m\beta\leftarrow m\alpha} & w_{m\beta\leftarrow f\beta} & w_{m\beta\leftarrow m\beta} \end{bmatrix} \tag{24}$$

where rows/columns of $\boldsymbol{W}$ denote destination/source classes while arrows in entry subscripts indicate transmission *directions* over each timestep. In this transmission process, a copy of the allele in a source class is expected to leave descendant copies in a destination class through *different* transmission pathways, which fall into *two* categories: the copy may persist through the survival of its bearer (this one contributes only when the source and destination classes are identical — i.e., the diagonal of $\boldsymbol{W}$), or be transmitted through reproduction to a newly recruited breeder in the destination class (when this class has available breeding vacancy). For transmissions through reproduction, a pathway depends on events that collectively define itself, such as whether the new recruit is locally born or immigrant, whether it is sired by a local or non-local male, whether the allele is transmitted maternally or paternally from the relevant class.

Each entry of $\boldsymbol{W}$ is derived by summing all transmission pathways through which a copy of the allele in source class $j$ is expected to leave descendant copies in destination class $i$ over one timestep (see diagrams illustrating transmission pathways below). More specifically, for each pathway linking $j$ and $i$, the probabilities of the associated events are *multiplied*, and such products are then *summed* across all the pathways that link $j$ and $i$ to define the entry $w_{i\leftarrow j}$. Thus, $\boldsymbol{W}$ gives the *expected* number of descendant copies in each destination class, produced by one copy currently in each source class, over one timestep. Note that in a particular local group of each type, the events defining those transmission pathways have their *realized* probabilistic outcomes. However, in the population, each grouptype is represented by *infinitely many* (genetically connected) groups, and when allele transmissions are *summarized* across groups of the same type, the *proportion* of transmissions following each pathway is represented by the *probability* of the pathway. Thus, $\boldsymbol{W}$ gives the population-level *deterministic expectations* instead of realized transmission outcomes of a particular local group.

Accordingly, let $\boldsymbol{z}_t = [z_{f\alpha},\ z_{m\alpha},\ z_{f\beta},\ z_{m\beta}]^\mathsf{T}$ denotes the vector of the expected allele copy abundances in the four sex-grouptype classes at timestep $t$, then the expected allele copy abundances at timestep $t+1$ are given by $\boldsymbol{z}_{t+1} = \boldsymbol{W}\boldsymbol{z}_t$ (a biologically meaningful initial vector $\boldsymbol{z}_0$ must have non-negative entries and at least one positive entry, as its entries give allele copy abundances). After reaching transmission stationarity, the allele copy abundances across the clases no longer change — i.e., $\boldsymbol{W}\boldsymbol{z}_t = \boldsymbol{z}_t$, where $\boldsymbol{z}_t$ is a *right* eigenvector of $\boldsymbol{W}$ (associated with the dominant eigenvalue $\lambda = 1$) that gives the *stationary distribution* of the allele copy abundances across the classes under these forward allele copy transmissions. While this iterative process describes how allele copy *abundances* change over timesteps, reproductive values describe how much a copy of the allele currently in each class is expected to contribute to its future descendant copies across the population, and are obtained from the dominant *left* eigenvector $\boldsymbol{v}$ of $\boldsymbol{W}$ — at allele transmission stationarity, $\boldsymbol{v}$ is also associated with the dominant eigenvalue $\lambda = 1$ of $\boldsymbol{W}$ [see (26) below].

The explicit expressions for the entries in (24), following the source-to-destination convention defined above, are derived as below.

$$w_{f\alpha\leftarrow f\alpha} = (1-\mu_f) + \frac{u\mu_f n_{f\alpha}}{2un_{f\alpha}}\left[\pi_{f\alpha} + (1-\pi_{f\alpha})\frac{un_{f\alpha}}{\bar{n}_f}\right] \tag{25.1}$$

$$w_{f\alpha\leftarrow m\alpha} = \frac{u\mu_f n_{f\alpha}}{2un_{m\alpha}}\left\{\pi_{f\alpha}\left[m_\alpha+(1-m_\alpha)\frac{un_{m\alpha}}{\bar{n}_m}\right] + (1-\pi_{f\alpha})\left[\frac{un_{f\alpha}}{\bar{n}_f}\left[m_\alpha+(1-m_\alpha)\frac{un_{m\alpha}}{\bar{n}_m}\right] + \left(1-\frac{un_{f\alpha}}{\bar{n}_f}\right)(1-m_\beta)\frac{un_{m\alpha}}{\bar{n}_m}\right]\right\} \tag{25.2}$$

$$w_{f\alpha\leftarrow f\beta} = \frac{u\mu_f n_{f\alpha}}{2\,(1-u)\,n_{f\beta}}(1-\pi_{f\alpha})\left(1-\frac{un_{f\alpha}}{\bar{n}_f}\right) \tag{25.3}$$

$$w_{f\alpha\leftarrow m\beta} = \frac{u\mu_f n_{f\alpha}}{2\,(1-u)\,n_{m\beta}}\left\{\pi_{f\alpha}(1-m_\alpha)\left(1-\frac{un_{m\alpha}}{\bar{n}_m}\right) + (1-\pi_{f\alpha})\left[\frac{un_{f\alpha}}{\bar{n}_f}(1-m_\alpha)\left(1-\frac{un_{m\alpha}}{\bar{n}_m}\right) + \left(1-\frac{un_{f\alpha}}{\bar{n}_f}\right)\left[m_\beta+(1-m_\beta)\left(1-\frac{un_{m\alpha}}{\bar{n}_m}\right)\right]\right]\right\} \tag{25.4}$$

$$w_{m\alpha\leftarrow f\alpha} = \frac{u\mu_m n_{m\alpha}}{2un_{f\alpha}}\left[\pi_{m\alpha} + (1-\pi_{m\alpha})\frac{un_{f\alpha}}{\bar{n}_f}\right] \tag{25.5}$$

$$w_{m\alpha\leftarrow m\alpha} = (1-\mu_m) + \frac{u\mu_m n_{m\alpha}}{2un_{m\alpha}}\left\{\pi_{m\alpha}\left[m_\alpha+(1-m_\alpha)\frac{un_{m\alpha}}{\bar{n}_m}\right] + (1-\pi_{m\alpha})\left[\frac{un_{f\alpha}}{\bar{n}_f}\left[m_\alpha+(1-m_\alpha)\frac{un_{m\alpha}}{\bar{n}_m}\right] + \left(1-\frac{un_{f\alpha}}{\bar{n}_f}\right)(1-m_\beta)\frac{un_{m\alpha}}{\bar{n}_m}\right]\right\} \tag{25.6}$$

$$w_{m\alpha\leftarrow f\beta} = \frac{u\mu_m n_{m\alpha}}{2\,(1-u)\,n_{f\beta}}(1-\pi_{m\alpha})\left(1-\frac{un_{f\alpha}}{\bar{n}_f}\right) \tag{25.7}$$

$$w_{m\alpha\leftarrow m\beta} = \frac{u\mu_m n_{m\alpha}}{2\,(1-u)\,n_{m\beta}}\left\{\pi_{m\alpha}(1-m_\alpha)\left(1-\frac{un_{m\alpha}}{\bar{n}_m}\right) + (1-\pi_{m\alpha})\left[\frac{un_{f\alpha}}{\bar{n}_f}\left[(1-m_\alpha)\left(1-\frac{un_{m\alpha}}{\bar{n}_m}\right)\right] + \left(1-\frac{un_{f\alpha}}{\bar{n}_f}\right)\left[m_\beta+(1-m_\beta)\left(1-\frac{un_{m\alpha}}{\bar{n}_m}\right)\right]\right]\right\} \tag{25.8}$$

$$w_{f\beta\leftarrow f\alpha} = \frac{(1-u)\,\mu_f n_{f\beta}}{2un_{f\alpha}}\left(1-\pi_{f\beta}\right)\frac{un_{f\alpha}}{\bar{n}_f} \tag{25.9}$$

$$w_{f\beta\leftarrow m\alpha} = \frac{(1-u)\,\mu_f n_{f\beta}}{2un_{m\alpha}}\left[\pi_{f\beta}\left(1-m_\beta\right)\frac{un_{m\alpha}}{\bar{n}_m} + \left(1-\pi_{f\beta}\right)\left[\frac{un_{f\alpha}}{\bar{n}_f}\left[m_\alpha + \left(1-m_\alpha\right)\frac{un_{m\alpha}}{\bar{n}_m}\right] + \left(1-\frac{un_{f\alpha}}{\bar{n}_f}\right)\left(1-m_\beta\right)\frac{un_{m\alpha}}{\bar{n}_m}\right]\right] \tag{25.10}$$

$$w_{f\beta\leftarrow f\beta} = \left(1-\mu_f\right) + \frac{(1-u)\,\mu_f n_{f\beta}}{2\,(1-u)\,n_{f\beta}}\left[\pi_{f\beta} + \left(1-\pi_{f\beta}\right)\left(1-\frac{un_{f\alpha}}{\bar{n}_f}\right)\right] \tag{25.11}$$

$$w_{f\beta\leftarrow m\beta} = \frac{(1-u)\,\mu_f n_{f\beta}}{2\,(1-u)\,n_{m\beta}}\left\{\pi_{f\beta}\left[m_\beta + \left(1-m_\beta\right)\left(1-\frac{un_{m\alpha}}{\bar{n}_m}\right)\right] + \left(1-\pi_{f\beta}\right)\left[\frac{un_{f\alpha}}{\bar{n}_f}\left(1-m_\alpha\right)\left(1-\frac{un_{m\alpha}}{\bar{n}_m}\right) + \left(1-\frac{un_{f\alpha}}{\bar{n}_f}\right)\left[m_\beta + \left(1-m_\beta\right)\left(1-\frac{un_{m\alpha}}{\bar{n}_m}\right)\right]\right]\right\} \tag{25.12}$$

$$w_{m\beta\leftarrow f\alpha} = \frac{(1-u)\,\mu_m n_{m\beta}}{2un_{f\alpha}}\left(1-\pi_{m\beta}\right)\frac{un_{f\alpha}}{\bar{n}_f} \tag{25.13}$$

$$w_{m\beta\leftarrow m\alpha} = \frac{(1-u)\,\mu_m n_{m\beta}}{2un_{m\alpha}}\left[\pi_{m\beta}\left(1-m_\beta\right)\frac{un_{m\alpha}}{\bar{n}_m} + \left(1-\pi_{m\beta}\right)\left[\frac{un_{f\alpha}}{\bar{n}_f}\left[m_\alpha + \left(1-m_\alpha\right)\frac{un_{m\alpha}}{\bar{n}_m}\right] + \left(1-\frac{un_{f\alpha}}{\bar{n}_f}\right)\left(1-m_\beta\right)\frac{un_{m\alpha}}{\bar{n}_m}\right]\right] \tag{25.14}$$

$$w_{m\beta\leftarrow f\beta} = \frac{(1-u)\,\mu_m n_{m\beta}}{2\,(1-u)\,n_{f\beta}}\left[\pi_{m\beta} + \left(1-\pi_{m\beta}\right)\left(1-\frac{un_{f\alpha}}{\bar{n}_f}\right)\right] \tag{25.15}$$

$$w_{m\beta\leftarrow m\beta} = \left(1-\mu_m\right) + \frac{(1-u)\,\mu_m n_{m\beta}}{2\,(1-u)\,n_{m\beta}}\left[\pi_{m\beta}\left[m_\beta + \left(1-m_\beta\right)\left(1-\frac{un_{m\alpha}}{\bar{n}_m}\right)\right] + \left(1-\pi_{m\beta}\right)\left[\frac{un_{f\alpha}}{\bar{n}_f}\left(1-m_\alpha\right)\left(1-\frac{un_{m\alpha}}{\bar{n}_m}\right) + \left(1-\frac{un_{f\alpha}}{\bar{n}_f}\right)\left[m_\beta + \left(1-m_\beta\right)\left(1-\frac{un_{m\alpha}}{\bar{n}_m}\right)\right]\right]\right] \tag{25.16}$$

For example, the flowchart below illustrates how equation (25.13), which accounts for the expected number of the allele copies transmitted (maternally) from a *female* in an $\alpha$-type group (or $f\alpha$ for simplicity) to a *male* in a $\beta$-type group (or $m\beta$ for simplicity), is derived (where nodes outlining all the *relevant* pathways of transmissions between the given classes are highlighted):

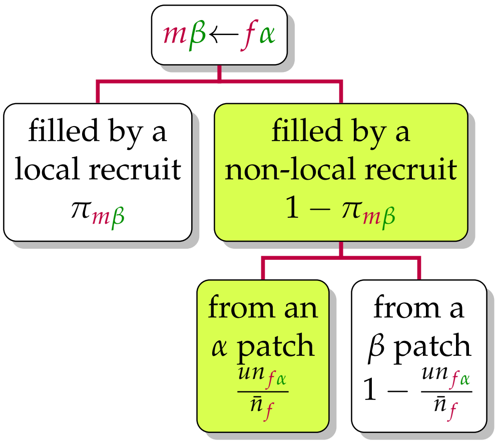

In (25.13) above, $\frac{(1-u)\mu_m n_{m\beta}}{2un_{f\alpha}}$ quantifies the per-allele rate at which *male* breeding vacancies (allele 'slots') in $\beta$-type group are *potentially available* to be 'filled' by alleles originating from *females* in $\alpha$-type groups (via these females' dispersing sons who win the 'fair-lottery' competition after arriving at $\beta$-type groups), where the numerator summarizes the total number of male allele 'slots' *available* across $\beta$-type groups (which are created/released by the deceased males in $\beta$-type groups — each death releases a breeding vacancy and provides one allele 'slot'), while the denominator summarizes the total number of (competing) alleles produced by females across $\alpha$-type groups (the factor of 2 accounts for diploidy — each female transmits either of her pair of alleles to each of its offspring maternally). Here, as we focus on the specific transmission $m\beta \leftarrow f\alpha$, only the pathway that a (released) male allele 'slot' in a $\beta$-type group is 'filled' by a non-local (or dispersed) male offspring (with probability $1-\pi_{m\beta}$) originating from an $\alpha$-type group (with probability $\frac{un_{f\alpha}}{\bar{n}_f}$) is relevant and thus considered in (25.13).

For another (more complicated) example, following the same biological reasoning, $w_{m\beta\leftarrow m\beta}$ [i.e., equation (25.16)] can be derived following the flowchart below (highlighted nodes outline all the relevant pathways of transmissions from $m\beta$ to $m\beta$ — in this case it involves not only reproduction, but also own survival):

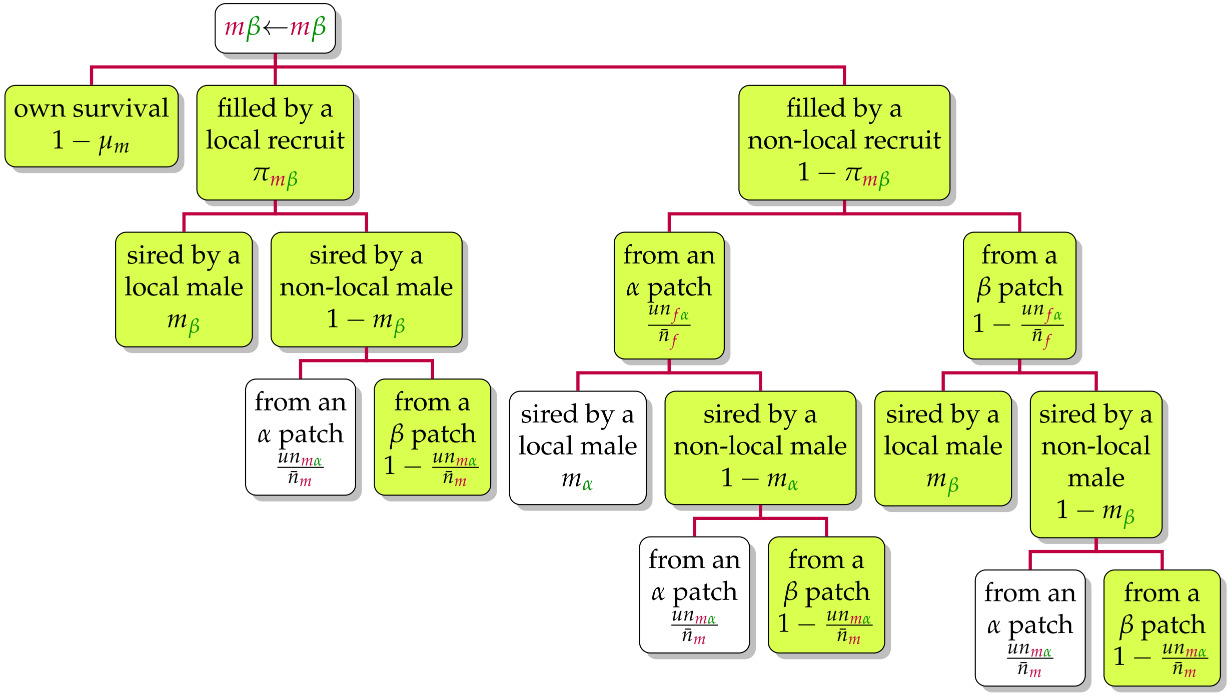

Likewise, by explicitly consider all the pathways under which allele copies can be transimitted from a given sex-grouptype class to another, we can derive the rest of the expressions given above for the matrix $\boldsymbol{W}$.

With entries of $\boldsymbol{W}$ explicitly expressed in (25.1) – (25.16), we can then derive the *sex-grouptype-specific* reproductive values of the allele, by solving

$$\boldsymbol{v}\boldsymbol{W} = \boldsymbol{v} \tag{26}$$

for $\boldsymbol{v}$. Biologically, $\boldsymbol{v}$ gives the expected relative contribution of a copy of the allele currently carried by a breeder in each sex-grouptype class to its future descendant copies (a higher value for a class means a copy currently in that class is expected to leave relatively more descendant copies in the population in the long run). We denote class-specific reproductive values as

$$\boldsymbol{v} = \begin{bmatrix} v_{f\alpha} & v_{m\alpha} & v_{f\beta} & v_{m\beta} \end{bmatrix} \tag{27}$$

As the absolute scale of $\boldsymbol{v}$ from (26) is arbitrary, for interpretability and consistency of our analysis, we normalize it as

$$\begin{aligned} \boldsymbol{v}' &= \begin{bmatrix} \frac{v_{f\alpha}}{\bar{v}} & \frac{v_{m\alpha}}{\bar{v}} & \frac{v_{f\beta}}{\bar{v}} & \frac{v_{m\beta}}{\bar{v}} \end{bmatrix} \\ &= \begin{bmatrix} v'_{f\alpha} & v'_{m\alpha} & v'_{f\beta} & v'_{m\beta} \end{bmatrix} \end{aligned} \tag{28}$$

where

$$\bar{v} = v_{f\alpha}\frac{un_{f\alpha}}{\bar{n}} + v_{m\alpha}\frac{un_{m\alpha}}{\bar{n}} + v_{f\beta}\frac{(1-u)n_{f\beta}}{\bar{n}} + v_{m\beta}\frac{(1-u)n_{m\beta}}{\bar{n}}$$

is the average reproductive value (i.e., sex-grouptype-specific reproductive values weighted by the relative abundances of breeders in each sex-grouptype class), and

$$\bar{n} = u(n_{f\alpha} + n_{m\alpha}) + (1-u)(n_{f\beta} + n_{m\beta})$$

is the (grouptype-frequency-weighted) average group size of the population. This normalization preserves the *relative* reproductive values among the classes while setting the breeder-abundance-weighted population average to 1 (thus a value above/below one indicates that a copy of the allele currently in the associated class has an expected above-/below-average contribution to the future descendant copies of the allele in the population).

##### 3.3 Fitness of an Allele for Helping/Harming

With reproductive values in (28) for breeders in any given sex-grouptype class, we can then derive the expressions (i.e., $w_{f\alpha}$, $w_{m\alpha}$, $w_{f\beta}$ and $w_{f\beta}$ below) capturing the fitness of the allele bore by a focal breeder in each of the sex-grouptype class — in terms of the expected number of allele copies the breeder transmits at each timestep at the population demographic equilibrium (given both its own fecundity and survival performances and those of other breeders).

Generally, fitness is expressed as a function of (I) the sex-grouptype-specific reproductive values, (II) the mortality and fecundity of the focal breeder itself, (III) the average mortality and fecundity of its groupmates, and (IV) the average mortality and fecundity of the breeders in the population. Following Ellis et al. (2022), these mortality and fecundity are realized at three social organizational levels: (I) those for the focal breeder at the individual level (i.e., own fitness performances), (II) those for the breeders in a given sex-grouptype class (i.e., the average fitness performances of breeders in the class — either including or excluding the focal breeder, depending on the specific contexts captured in equations 29 - 32 below), and (III) those for the breeders across the population (i.e., the average population-level fitness performances of breeders).

As before, we use $\mu$ and $p$ to denote mortality and fecundity, respectively, and '$f$', '$m$', '$\alpha$', and '$\beta$' in subscripts to indicate the sexes and grouptypes. To distinguish fecundity and mortality at different levels, we introduce subscripts '$i$' and '$g$' to indicate whether they are for a focal individual or the group of the focal breeder, respectively, while the average mortality and fecundity across the population are only sex-specific and thus indicated only by '$f$' or '$m$' in the subscripts. Besides, we use asterisks as superscripts to indicate the contexts where the group-level fecundity and mortality are defined *without* considering those of the focal breeder. Then, the sex-grouptype-specific fitness functions can be expressed as below — by exhaustively enumerating every possible pathway the mutant allele underlying the social behaviour can take to the next timestep (via survival and reproduction).

$$
\begin{aligned}
w_{f\alpha} = {} & v'_{f\alpha}\left\{(1-\mu_{if\alpha}) + \frac{(1-d_f)xp_{if\alpha}\left[\mu_{if\alpha}+(n_{f\alpha}-1)\mu^*_{gf\alpha}\right]}{2\left[(1-d_f)xp_{if\alpha}+(1-d_f)(n_{f\alpha}-1)xp^*_{gf\alpha}+d_f\bar{n}_f xp_f\right]} + \frac{d_f xp_{if\alpha}u\mu_f n_{f\alpha}}{2\left[(1-d_f)xp_f n_{f\alpha}+d_f xp_f\bar{n}_f\right]}\right\}+ \\
& v'_{m\alpha}\left\{\frac{(1-d_m)(1-x)p_{if\alpha}\mu_{gm\alpha}n_{m\alpha}}{2\left[(1-d_m)(1-x)p_{if\alpha}+(1-d_m)(n_{f\alpha}-1)(1-x)p^*_{gf\alpha}+d_m\bar{n}_f(1-x)p_f\right]} + \frac{d_m(1-x)p_{if\alpha}u\mu_m n_{m\alpha}}{2\left[(1-d_m)(1-x)p_f n_{f\alpha}+d_m(1-x)p_f\bar{n}_f\right]}\right\}+ \\
& v'_{f\beta}\left\{\frac{d_f xp_{if\alpha}(1-u)\mu_f n_{f\beta}}{2\left[(1-d_f)xp_f n_{f\beta}+d_f xp_f\bar{n}_f\right]}\right\}+ \\
& v'_{m\beta}\left\{\frac{d_m(1-x)p_{if\alpha}(1-u)\mu_m n_{m\beta}}{2\left[(1-d_m)(1-x)p_f n_{f\beta}+d_m(1-x)p_f\bar{n}_f\right]}\right\}
\end{aligned}
\tag{29}
$$

$$
\begin{aligned}
w_{f\beta} = {} & v'_{f\alpha}\left\{\frac{d_f xp_{if\beta}u\mu_f n_{f\alpha}}{2\left[(1-d_f)xp_f n_{f\alpha}+d_f xp_f\bar{n}_f\right]}\right\}+ \\
& v'_{m\alpha}\left\{\frac{d_m(1-x)p_{if\beta}u\mu_m n_{m\alpha}}{2\left[(1-d_m)(1-x)p_f n_{f\alpha}+d_m(1-x)p_f\bar{n}_f\right]}\right\}+ \\
& v'_{f\beta}\left\{(1-\mu_{if\beta}) + \frac{(1-d_f)xp_{if\beta}\left[\mu_{if\beta}+(n_{f\beta}-1)\mu^*_{gf\beta}\right]}{2\left[(1-d_f)xp_{if\beta}+(1-d_f)(n_{f\beta}-1)xp^*_{gf\beta}+d_f\bar{n}_f xp_f\right]} + \frac{d_f xp_{if\beta}(1-u)\mu_f n_{f\beta}}{2\left[(1-d_f)xp_f n_{f\beta}+d_f xp_f\bar{n}_f\right]}\right\}+ \\
& v'_{m\beta}\left\{\frac{(1-d_m)(1-x)p_{if\beta}\mu_{gm\beta}n_{m\beta}}{2\left[(1-d_m)(1-x)p_{if\beta}+(1-d_m)(n_{f\beta}-1)(1-x)p^*_{gf\beta}+d_m\bar{n}_f(1-x)p_f\right]} + \frac{d_m(1-x)p_{if\beta}(1-u)\mu_m n_{m\beta}}{2\left[(1-d_m)(1-x)p_f n_{f\beta}+d_m(1-x)p_f\bar{n}_f\right]}\right\}
\end{aligned}
\tag{30}
$$

$$
\begin{aligned}
w_{m\alpha} = v'_{f\alpha} \Bigg\{ & \Bigg[ \frac{(1-d_f)xp_{gf\alpha}n_{f\alpha}\mu_{gf\alpha}n_{f\alpha}}{2\left[(1-d_f)xp_{gf\alpha}n_{f\alpha}+d_fxp_f\bar{n}_f\right]} + \frac{d_fxp_{gf\alpha}n_{f\alpha}u\mu_fn_{f\alpha}}{2\left[(1-d_f)xp_fn_{f\alpha}+d_fxp_f\bar{n}_f\right]} \Bigg] \frac{mp_{im\alpha}}{mp_{im\alpha}+m(n_{m\alpha}-1)p^*_{gm\alpha}+(1-m)\bar{n}_mp_m} + \\
& \frac{(1-m)p_{im\alpha}u}{[mp_mn_{m\alpha}+(1-m)p_m\bar{n}_m]} \Big[ \frac{(1-d_f)xp_fn_{f\alpha}\mu_fn_{f\alpha}}{2\left[(1-d_f)xp_fn_{f\alpha}+d_fxp_f\bar{n}_f\right]} + \frac{d_fxp_fn_{f\alpha}u\mu_fn_{f\alpha}}{2\left[(1-d_f)xp_fn_{f\alpha}+d_fxp_f\bar{n}_f\right]} \Big] + \\
& \frac{(1-m)p_{im\alpha}(1-u)}{[mp_mn_{m\beta}+(1-m)p_m\bar{n}_m]} \frac{d_fxp_fn_{f\beta}u\mu_fn_{f\alpha}}{2\left[(1-d_f)xp_fn_{f\alpha}+d_fxp_f\bar{n}_f\right]} \Bigg\} + \\
v'_{m\alpha} \Bigg\{ & (1-\mu_{im\alpha}) + \\
& \Bigg[ \frac{(1-d_m)(1-x)p_{gf\alpha}n_{f\alpha}\left[\mu_{im\alpha}+(n_{m\alpha}-1)\mu^*_{gm\alpha}\right]}{2\left[(1-d_m)(1-x)p_{gf\alpha}n_{f\alpha}+d_m(1-x)p_f\bar{n}_f\right]} + \frac{d_m(1-x)p_{gf\alpha}n_{f\alpha}u\mu_mn_{m\alpha}}{2\left[(1-d_m)(1-x)p_fn_{f\alpha}+d_m(1-x)p_f\bar{n}_f\right]} \Bigg] \frac{mp_{im\alpha}}{mp_{im\alpha}+m(n_{m\alpha}-1)p^*_{gm\alpha}+(1-m)\bar{n}_mp_m} + \\
& \frac{(1-m)p_{im\alpha}u}{[mp_mn_{m\alpha}+(1-m)p_m\bar{n}_m]} \Big[ \frac{(1-d_m)(1-x)p_fn_{f\alpha}\mu_mn_{m\alpha}}{2\left[(1-d_m)(1-x)p_fn_{f\alpha}+d_m(1-x)p_f\bar{n}_f\right]} + \frac{d_m(1-x)p_fn_{f\alpha}u\mu_mn_{m\alpha}}{2\left[(1-d_m)(1-x)p_fn_{f\alpha}+d_m(1-x)p_f\bar{n}_f\right]} \Big] + \\
& \frac{(1-m)p_{im\alpha}(1-u)}{[mp_mn_{m\beta}+(1-m)p_m\bar{n}_m]} \frac{d_m(1-x)p_fn_{f\beta}u\mu_mn_{m\alpha}}{2\left[(1-d_m)(1-x)p_fn_{f\alpha}+d_m(1-x)p_f\bar{n}_f\right]} \Bigg\} + \\
v'_{f\beta} \Bigg\{ & \frac{d_fxp_{gf\alpha}n_{f\alpha}(1-u)\mu_fn_{f\beta}}{2\left[(1-d_f)xp_fn_{f\beta}+d_fxp_f\bar{n}_f\right]} \frac{mp_{im\alpha}}{mp_{im\alpha}+m(n_{m\alpha}-1)p^*_{gm\alpha}+(1-m)\bar{n}_mp_m} + \\
& \frac{(1-m)p_{im\alpha}u}{[mp_mn_{m\alpha}+(1-m)p_m\bar{n}_m]} \frac{d_fxp_fn_{f\alpha}(1-u)\mu_fn_{f\beta}}{2\left[(1-d_f)xp_fn_{f\beta}+d_fxp_f\bar{n}_f\right]} + \\
& \frac{(1-m)p_{im\alpha}(1-u)}{[mp_mn_{m\beta}+(1-m)p_m\bar{n}_m]} \Big[ \frac{(1-d_f)xp_fn_{f\beta}\mu_fn_{f\beta}}{2\left[(1-d_f)xp_fn_{f\beta}+d_fxp_f\bar{n}_f\right]} + \frac{d_fxp_fn_{f\beta}(1-u)\mu_fn_{f\beta}}{2\left[(1-d_f)xp_fn_{f\beta}+d_fxp_f\bar{n}_f\right]} \Big] \Bigg\} + \\
v'_{m\beta} \Bigg\{ & \frac{d_m(1-x)p_{gf\alpha}n_{f\alpha}(1-u)\mu_mn_{m\beta}}{2\left[(1-d_m)(1-x)p_fn_{f\beta}+d_m(1-x)p_f\bar{n}_f\right]} \frac{mp_{im\alpha}}{mp_{im\alpha}+m(n_{m\alpha}-1)p^*_{gm\alpha}+(1-m)\bar{n}_mp_m} + \\
& \frac{(1-m)p_{im\alpha}u}{[mp_mn_{m\alpha}+(1-m)p_m\bar{n}_m]} \frac{d_m(1-x)p_fn_{f\alpha}(1-u)\mu_mn_{m\beta}}{2\left[(1-d_m)(1-x)p_fn_{f\beta}+d_m(1-x)p_f\bar{n}_f\right]} + \\
& \frac{(1-m)p_{im\alpha}(1-u)}{[mp_mn_{m\beta}+(1-m)p_m\bar{n}_m]} \Big[ \frac{(1-d_m)(1-x)p_fn_{f\beta}\mu_mn_{m\beta}}{2\left[(1-d_m)(1-x)p_fn_{f\beta}+d_m(1-x)p_f\bar{n}_f\right]} + \frac{d_m(1-x)p_fn_{f\beta}(1-u)\mu_mn_{m\beta}}{2\left[(1-d_m)(1-x)p_fn_{f\beta}+d_m(1-x)p_f\bar{n}_f\right]} \Big] \Bigg\}
\end{aligned}
\tag{31}
$$

$$
\begin{aligned}
w_{m\beta} = {} & v'_{f\alpha}\Bigg\{ \frac{d_f x p_{gf\beta} n_{f\beta} u \mu_f n_{f\alpha}}{2\left[(1-d_f)xp_f n_{f\alpha} + d_f x p_f \bar{n}_f\right]} \frac{m p_{im\beta}}{m p_{im\beta} + m(n_{m\beta}-1)p^*_{gm\beta} + (1-m)\bar{n}_m p_m} + \\
& \frac{(1-m)p_{im\beta}u}{[m p_m n_{m\alpha} + (1-m)p_m \bar{n}_m]}\Big[\frac{(1-d_f)xp_f n_{f\alpha}\mu_f n_{f\alpha}}{2\left[(1-d_f)xp_f n_{f\alpha} + d_f x p_f \bar{n}_f\right]} + \frac{d_f x p_f n_{f\alpha} u \mu_f n_{f\alpha}}{2\left[(1-d_f)xp_f n_{f\alpha} + d_f x p_f \bar{n}_f\right]}\Big] + \\
& \frac{(1-m)p_{im\beta}(1-u)}{[m p_m n_{m\beta} + (1-m)p_m \bar{n}_m]} \frac{d_f x p_f n_{f\beta} u \mu_f n_{f\alpha}}{2\left[(1-d_f)xp_f n_{f\alpha} + d_f x p_f \bar{n}_f\right]}\Bigg\} + \\
& v'_{m\alpha}\Bigg\{ \frac{d_m(1-x) p_{gf\beta} n_{f\beta} u \mu_m n_{m\alpha}}{2\left[(1-d_m)(1-x)p_f n_{f\alpha} + d_m(1-x) p_f \bar{n}_f\right]} \frac{m p_{im\beta}}{m p_{im\beta} + m(n_{m\beta}-1)p^*_{gm\beta} + (1-m)\bar{n}_m p_m} + \\
& \frac{(1-m)p_{im\beta}u}{[m p_m n_{m\alpha} + (1-m)p_m \bar{n}_m]}\Big[\frac{(1-d_m)(1-x)p_f n_{f\alpha}\mu_m n_{m\alpha}}{2\left[(1-d_m)(1-x)p_f n_{f\alpha} + d_m(1-x) p_f \bar{n}_f\right]} + \frac{d_m(1-x) p_f n_{f\alpha} u \mu_m n_{m\alpha}}{2\left[(1-d_m)(1-x)p_f n_{f\alpha} + d_m(1-x) p_f \bar{n}_f\right]}\Big] + \\
& \frac{(1-m)p_{im\beta}(1-u)}{[m p_m n_{m\beta} + (1-m)p_m \bar{n}_m]} \frac{d_m(1-x) p_f n_{f\beta} u \mu_m n_{m\alpha}}{2\left[(1-d_m)(1-x)p_f n_{f\alpha} + d_m(1-x) p_f \bar{n}_f\right]}\Bigg\} + \\
& v'_{f\beta}\Bigg\{ \left[\frac{(1-d_f)xp_{gf\beta} n_{f\beta}\mu_{gf\beta} n_{f\beta}}{2\left[(1-d_f)xp_{gf\beta} n_{f\beta} + d_f x p_f \bar{n}_f\right]} + \frac{d_f x p_{gf\beta} n_{f\beta}(1-u)\mu_f n_{f\beta}}{2\left[(1-d_f)xp_f n_{f\beta} + d_f x p_f \bar{n}_f\right]}\right] \frac{m p_{im\beta}}{m p_{im\beta} + m(n_{m\beta}-1)p^*_{gm\beta} + (1-m)\bar{n}_m p_m} + \\
& \frac{(1-m)p_{im\beta}u}{[m p_m n_{m\alpha} + (1-m)p_m \bar{n}_m]} \frac{d_f x p_f n_{f\alpha}(1-u)\mu_f n_{f\beta}}{2\left[(1-d_f)xp_f n_{f\beta} + d_f x p_f \bar{n}_f\right]} + \\
& \frac{(1-m)p_{im\beta}(1-u)}{[m p_m n_{m\beta} + (1-m)p_m \bar{n}_m]}\Big[\frac{(1-d_f)xp_f n_{f\beta}\mu_f n_{f\beta}}{2\left[(1-d_f)xp_f n_{f\beta} + d_f x p_f \bar{n}_f\right]} + \frac{d_f x p_f n_{f\beta}(1-u)\mu_f n_{f\beta}}{2\left[(1-d_f)xp_f n_{f\beta} + d_f x p_f \bar{n}_f\right]}\Big]\Bigg\} + \\
& v'_{m\beta}\Bigg\{(1-\mu_{im\beta}) + \\
& \left[\frac{(1-d_m)(1-x)p_{gf\beta} n_{f\beta}\left[\mu_{im\beta} + (n_{m\beta}-1)\mu^*_{gm\beta}\right]}{2\left[(1-d_m)(1-x)p_{gf\beta} n_{f\beta} + d_m(1-x) p_f \bar{n}_f\right]} + \frac{d_m(1-x) p_{gf\beta} n_{f\beta}(1-u)\mu_m n_{m\beta}}{2\left[(1-d_m)(1-x)p_f n_{f\beta} + d_m(1-x) p_f \bar{n}_f\right]}\right] \frac{m p_{im\beta}}{m p_{im\beta} + m(n_{m\beta}-1)p^*_{gm\beta} + (1-m)\bar{n}_m p_m} + \\
& \frac{(1-m)p_{im\beta}u}{[m p_m n_{m\alpha} + (1-m)p_m \bar{n}_m]} \frac{d_m(1-x) p_f n_{f\alpha}(1-u)\mu_m n_{m\beta}}{2\left[(1-d_m)(1-x)p_f n_{f\beta} + d_m(1-x) p_f \bar{n}_f\right]} + \\
& \frac{(1-m)p_{im\beta}(1-u)}{[m p_m n_{m\beta} + (1-m)p_m \bar{n}_m]}\Big[\frac{(1-d_m)(1-x)p_f n_{f\beta}\mu_m n_{m\beta}}{2\left[(1-d_m)(1-x)p_f n_{f\beta} + d_m(1-x) p_f \bar{n}_f\right]} + \frac{d_m(1-x) p_f n_{f\beta}(1-u)\mu_m n_{m\beta}}{2\left[(1-d_m)(1-x)p_f n_{f\beta} + d_m(1-x) p_f \bar{n}_f\right]}\Big]\Bigg\}
\end{aligned} \tag{32}
$$

As we see, both the fractions of female (i.e., $x$) and male (i.e., $1 - x$) offspring cancel out in (29) – (32), thus offspring sex ratio does not affect fitness. Below, we illustrate how these fitness functions are derived for each sex-grouptype class. For simplicity, we focus on describing the derivation of (29) as an illustrative example; equations (30) – (32) are derived by explicitly accounting for the contributions of the possible pathways by which a focal breeder of given sex and grouptype transmits the allele, from one timestep to the next under population demographic equilibrium, through its own survival and/or the survival of its offspring in the population, following the same logic as in the derivation of (29).

Equation (29) captures the expected number of allele copies a female breeder in an $\alpha$-type group transmits at each timestep at the population demographic equilibrium. Each line on the right hand side (*rhs*) of the expression captures a pathway under which an expected amount of allele copies is accounted for. Specifically, the first line captures the expected amount allele copies attributed to the focal female's own sex-grouptype class (weighted by the expected reproductive value for that class). Here, the first term captures the expected number of allele copies bore by the focal female herself from one timestep to the next, via her own survival to the next timestep (she can only transmit a single copy of the allele if she survives to the next timestep). The second term captures the expected amount of allele transmitted to the focal female's own *daughters* who will stay at the focal's group at the next timestep (i.e., become an established breeder). Here, the numerator is the share of the 'allele slots' of her philopatric daughters, while the denominator is the total number of 'allele slots' available to all the *female* offspring in the group (i.e., native female offspring who stay and those female offspring expected to arrive at the group) to compete for (as we focus on diploid each breeding vacancy is associated with 2 'allele slots'). The third term is the expected amount of 'allele slots' the *daughters* of the focal female are expected to claim *elsewhere* in the population (these daughters are expected to disperse to other $\alpha$-type groups and win their breeding vacancies).

The second line captures the expected amount of allele copies attributed to the males in the focal female's group (weighted by the expected reproductive value for each male breeder in $\alpha$-type group). Here, the numerator in the first term captures the share of 'allele slots' the focal female's *philopatric sons* are expected to claim in the next timestep, while the denominator account for the expected total amount of 'allele slots' available for *male* offspring to claim (by competing with the philopatric male offspring in the group along with those from elsewhere in the population). The second term in this line accounts for the number of 'allele slots' the focal female's sons are expected to claim in other $\alpha$-type groups in the population (these sons of the focal female are expected to disperse and succeed in the fair lottery in other groups of the same type).

The third line captures the expected amount of allele copies attributed to the female offspring in $\beta$-type groups in the population (weighted by the expected reproductive value for each female breeder in $\beta$-type group). Here, the numerator captures the number of 'allele slots' the focal female's daughters are expected to share in $\beta$-type groups in the population (these daughters are expected to disperse to $\beta$-type groups and succeed in the fair lottery competition in these $\beta$-type groups).

The last line captures the expected amount of allele copies attributed to the male offspring in $\beta$-type groups in the population (weighted by the expected reproductive value for each male breeder in $\beta$-type group). Here, the numerator captures the amount of 'allele slots' the focal female's sons are expected to claim in $\beta$-type groups in the population (these sons are expected to disperse to $\beta$-type groups and succeed in the fair lottery competition in this type of groups).

##### 3.4 Selection on Helping/Harming

With the above fitness expressions, we can then evaluate the inclusive fitness effects of the behaviour underpinned by the allele in either grouptype — provided that it affects the fecundity or mortality of individuals rather than being neutral. We assume the behaviour imposes some total marginal fitness benefits or costs of magnitude $b$ (say $0 < |b| \ll 1$) to the local recipients of either sex, in terms of either fecundity (3.4.1 below) or mortality (3.4.2 below), where $b > 0$ implies helping while $b < 0$ indicates harming. We assume these benefits or costs are equally divided among targeted recipients, and that expressing the behaviour by a focal individual incurs a constant marginal fecundity or mortality cost of magnitude $c$ ($0 < c \ll 1$). Under these assumptions, we can derive the expressions accounting for the inclusive fitness outcomes of the behaviour by the focal, found in a given *sex*, *age* and *grouptype*, directed to groupmates (recipients), found in a given *sex*, by applying the neighbor-modulated inclussive fitness approach (Taylor and Frank, 1996) in these explicit contexts. For each of these expressions, we can find the critical value $b^*$ at which the behaviour would have *no* inclusive fitness impact (i.e., the behaviour is at the equilibrium condition), and use the critical ratio $c/b^*$ as a measure of both the *direction* and *strength* of selection on the behaviour — when $c/b^* > 0$, the more this (positive) ratio deviates from 0, the stronger the selective pressure for helping, while when $c/b^* < 0$, the more this (negative) ratio deviates from 0, the stronger the selective pressure for harming.

The expressions capturing the above-mentioned inclusive fitness outcomes of the behaviour are given in equations (33.1) – (36.4) below. Briefly, these expressions are derived by following the same biological reasoning and logical steps: we (I) begin with the relevant *direct* fitness functions from (29) – (32), (II) apply the marginal social behaviour (the actor pays a personal fecundity or mortality cost while providing a total fecundity or mortality benefit or cost equally divided among all recipients of the targeted sex), and then (III) calculate the total inclusive fitness effect as the sum of four major components — which, for clarity, we highlight below in distinct colours to visually map them to the corresponding terms appearing in each of the equations below:

- the direct fecundity/mortality cost to the actor,
- the feedback fecundity/mortality effect on the actor through changes in the 'excluding-self' group average of the actor's own sex — this component appears *only* when the targeted (or recipient) sex is the same as the actor's sex (i.e., same-sex interaction),
- the indirect fecundity/mortality effects on the recipients, incorporating exact finite-group corrections: when the actor and recipients are the *same sex*, each recipient's 'excluding-self' average is influenced by both the actor's personal cost and the benefits/costs received by the other recipients, while when the actor and recipients are *opposite sexes*, the actor's personal cost does not affect the recipients' 'excluding-self' averages (i.e., only the direct effect on each recipient and the feedback from the other recipients are included), and
- the spillover fecundity/mortality effects on individuals of the non-targeted sex (if exists), resulting from the net change in the inclusive group average of the targeted sex (this net change includes the actor's personal cost whenever the actor belongs to the targeted sex).

##### 3.4.1 The behaviour with fecundity consequences

When the *fecundity* benefits (or costs) by the behaviour are directed to other *females* at the fecundity cost of a focal *female* aged $a$ in an $\alpha$-type group (indicated by superscripts on the left hand side of the equations below), the inclusive fitness outcomes are captured by

$$F_{ff}^{\alpha a} = -c\frac{\partial w_{f\alpha}}{\partial p_{if\alpha}} + \frac{b}{n_{f\alpha}-1}\frac{\partial w_{f\alpha}}{\partial p_{gf\alpha}^{*}} + \left(n_{f\alpha}-1\right) r_{ff}^{\alpha a}\left[\frac{b}{n_{f\alpha}-1}\frac{\partial w_{f\alpha}}{\partial p_{if\alpha}} + \frac{1}{n_{f\alpha}-1}\left[\left(n_{f\alpha}-2\right)\frac{b}{n_{f\alpha}-1} - c\right]\frac{\partial w_{f\alpha}}{\partial p_{gf\alpha}^{*}}\right] + n_{m\alpha} r_{mf}^{\alpha a}\frac{b-c}{n_{f\alpha}}\frac{\partial w_{m\alpha}}{\partial p_{gf\alpha}} \tag{33.1}$$

where '$F$' indicates that the behaviour affects *fecundity* — its first and second subscripts indicate the *recipient* and *focal* sex, respectively. The context under which equation (33.1) is formulated is illustrated by Fig. A0.

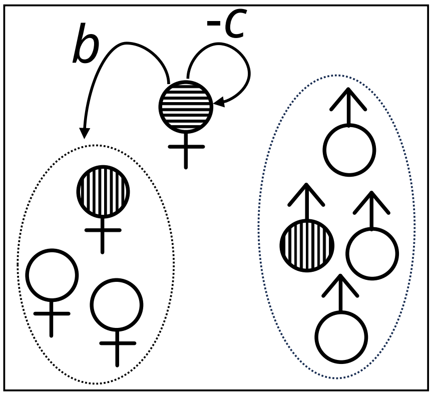

**Fig. A0** Schematic diagram illustrating the context under which equation (33.1) is formulated for an $\alpha$-type group (assuming 4 females and 4 males). Here, both the actor (shown with horizontal stripes) and the 3 recipients (in the dotted ellipse) are females while the non-targeted sex are the 4 males (in the dotted ellipse). From the perspective of the actor, her behavior confers a direct fecundity cost to herself, and a benefit (or cost) to these local recipients (distributed equally among them) — impacting the average fecundity of these recipient females also generates a fecundity feedback to the actor herself. From the perspective of a recipient female (shown with vertical stripes), she sees that (i) her own fecundity changes, and (ii) the average fecundity of other females (i.e., the actor and the other two female recipients) also changes. From the perspective of a male (shown with vertical stripes), his fecundity changes when the average fecundity of his female groupmates changes.

More specifically, the first line on the *rhs* of (33.1) quantifies the fecundity effects on the focal female herself. More specifically

- The first term ($-c\frac{\partial w_{f\alpha}}{\partial p_{if\alpha}}$) is the *direct personal cost*: the focal female reduces her own fecundity by a (tiny) magnitude of $c$.
- The second term ($\frac{b}{n_{f\alpha}-1}\frac{\partial w_{f\alpha}}{\partial p^{*}_{gf\alpha}}$) is the *neighbor-modulated feedback* to the focal: by making the other females more (or less) fecund, the focal female changes the competitive environment her own offspring will experience. In the model, $\frac{\partial w_{f\alpha}}{\partial p^{*}_{gf\alpha}} < 0$, so when $b > 0$ (helping) the feedback is *negative* (i.e., more competing offspring from other non-focal females makes it harder for the focal's own offspring to claim available breeding vacancies); when $b < 0$ (harming) the feedback is *positive* (i.e., fewer competitors the focal's offspring will experience). This term does *not* change the focal's own fecundity — her $p_{if\alpha}$ remains unchanged; only the *effectiveness* of her reproduction changes via increased/decreased offspring competition.

The second line on the *rhs* of (33.1) summarizes the indirect fecundity effects of the focal's behaviour on the other $n_{f\alpha} - 1$ recipient females, weighted by $r^{\alpha a}_{ff}$. From the perspective of a *recipient* female, her fecundity changes in two ways: (a) *direct benefit or cost to herself*: her own fecundity increases or decreases by a magnitude of $\frac{b}{n_{f\alpha}-1}$ (if the focal helps $b > 0$ or $b < 0$ if it harms), where the marginal value is $\frac{\partial w_{f\alpha}}{\partial p_{if\alpha}}$ (which has the same functional form as for the actor), and (b) *fecundity feedback to her via the changed group environment*: she sees that the actor paid a cost of magnitude $c$ (i.e., the actor has slightly lower fecundity), and the other $n_{f\alpha} - 2$ recipients each gained a benefit or cost of magnitude $\frac{b}{n_{f\alpha}-1}$. Thus, she sees that the 'excluding-self' average fecundity changes by a magnitude of $\frac{1}{n_{f\alpha}-1}\left[\left(n_{f\alpha}-2\right)\frac{b}{n_{f\alpha}-1} - c\right]$, which is derived as a finite-group correction.

The third line on the *rhs* of (33.1) captures the *indirect spillover effects* on the $n_{m\alpha}$ males, weighted by $r^{\alpha a}_{mf}$. Although the behaviour directly targets only female fecundity in this context, males' reproductive success rides on female productivity (i.e., *female demographic dominance*). The magnitude of *net* change in average *female* fecundity (*including* the actor) is $\frac{b-c}{n_{f\alpha}}$ (i.e., total benefit or cost of magnitude $b$ to other females and a direct cost of magnitude $c$ to actor, divided by the total number of females $n_{f\alpha}$ for an $\alpha$-type group). Therefore, from a male's perspective, he experiences this average change in female fecundity, which affects his own fitness via $\frac{\partial w_{m\alpha}}{\partial p_{gf\alpha}} > 0$. That is, the focal female's social behaviour indirectly helps or harms her *male* groupmates through changing the overall reproductive output of the *females* in the group.

The rest of the expressions for the behaviour with *fecundity* effect by a focal individual with given *sex* and aged $a$ in an $\alpha$-type group thus are:

$$F_{mf}^{\alpha a} = -c\frac{\partial w_{f\alpha}}{\partial p_{if\alpha}} + \left(n_{f\alpha}-1\right) r_{ff}^{\alpha a}\frac{-c}{n_{f\alpha}-1}\frac{\partial w_{f\alpha}}{\partial p_{gf\alpha}^{*}} + n_{m\alpha} r_{mf}^{\alpha a}\left[\frac{b}{n_{m\alpha}}\frac{\partial w_{m\alpha}}{\partial p_{im\alpha}} + \frac{b}{n_{m\alpha}}\frac{\partial w_{m\alpha}}{\partial p_{gm\alpha}^{*}} + \frac{-c}{n_{f\alpha}}\frac{\partial w_{m\alpha}}{\partial p_{gf\alpha}}\right] \tag{33.2}$$

$$F_{fm}^{\alpha a} = -c\frac{\partial w_{m\alpha}}{\partial p_{im\alpha}} + \frac{b}{n_{f\alpha}}\frac{\partial w_{m\alpha}}{\partial p_{gf\alpha}} + n_{f\alpha} r_{fm}^{\alpha a}\left[\frac{b}{n_{f\alpha}}\frac{\partial w_{f\alpha}}{\partial p_{if\alpha}} + \frac{b}{n_{f\alpha}}\frac{\partial w_{f\alpha}}{\partial p_{gf\alpha}^{*}}\right] + \left(n_{m\alpha}-1\right) r_{mm}^{\alpha a}\left(\frac{-c}{n_{m\alpha}-1}\frac{\partial w_{m\alpha}}{\partial p_{gm\alpha}^{*}} + \frac{b}{n_{f\alpha}}\frac{\partial w_{m\alpha}}{\partial p_{gf\alpha}}\right) \tag{33.3}$$

$$F_{mm}^{\alpha a} = -c\frac{\partial w_{m\alpha}}{\partial p_{im\alpha}} + \frac{b}{n_{m\alpha}-1}\frac{\partial w_{m\alpha}}{\partial p_{gm\alpha}^{*}} + \left(n_{m\alpha}-1\right) r_{mm}^{\alpha a}\left[\frac{b}{n_{m\alpha}-1}\frac{\partial w_{m\alpha}}{\partial p_{im\alpha}} + \frac{1}{n_{m\alpha}-1}\left[\left(n_{m\alpha}-2\right)\frac{b}{n_{m\alpha}-1} - c\right]\frac{\partial w_{m\alpha}}{\partial p_{gm\alpha}^{*}}\right] \tag{33.4}$$

Analogously, in a $\beta$-type group, the expressions for the behaviour with *fecundity* effect are

$$
\begin{aligned}
F_{ff}^{\beta a} = & -c\frac{\partial w_{f\beta}}{\partial p_{if\beta}} + \frac{b}{n_{f\beta}-1}\frac{\partial w_{f\beta}}{\partial p^{*}_{gf\beta}} + \\
& \left(n_{f\beta}-1\right) r_{ff}^{\beta a}\left[\frac{b}{n_{f\beta}-1}\frac{\partial w_{f\beta}}{\partial p_{if\beta}} + \frac{1}{n_{f\beta}-1}\left[\left(n_{f\beta}-2\right)\frac{b}{n_{f\beta}-1} - c\right]\frac{\partial w_{f\beta}}{\partial p^{*}_{gf\beta}}\right] + \\
& n_{m\beta} r_{mf}^{\beta a}\frac{b-c}{n_{f\beta}}\frac{\partial w_{m\beta}}{\partial p_{gf\beta}}
\end{aligned} \tag{34.1}
$$

$$
\begin{aligned}
F_{mf}^{\beta a} = & -c\frac{\partial w_{f\beta}}{\partial p_{if\beta}} + \\
& \left(n_{f\beta}-1\right) r_{ff}^{\beta a}\frac{-c}{n_{f\beta}-1}\frac{\partial w_{f\beta}}{\partial p^{*}_{gf\beta}} + \\
& n_{m\beta} r_{mf}^{\beta a}\left[\frac{b}{n_{m\beta}}\frac{\partial w_{m\beta}}{\partial p_{im\beta}} + \frac{b}{n_{m\beta}}\frac{\partial w_{m\beta}}{\partial p^{*}_{gm\beta}} + \frac{-c}{n_{f\beta}}\frac{\partial w_{m\beta}}{\partial p_{gf\beta}}\right]
\end{aligned} \tag{34.2}
$$

$$
\begin{aligned}
F_{fm}^{\beta a} = & -c\frac{\partial w_{m\beta}}{\partial p_{im\beta}} + \frac{b}{n_{f\beta}}\frac{\partial w_{m\beta}}{\partial p_{gf\beta}} + \\
& n_{f\beta} r_{fm}^{\beta a}\left[\frac{b}{n_{f\beta}}\frac{\partial w_{f\beta}}{\partial p_{if\beta}} + \frac{b}{n_{f\beta}}\frac{\partial w_{f\beta}}{\partial p^{*}_{gf\beta}}\right] + \\
& \left(n_{m\beta}-1\right) r_{mm}^{\beta a}\left(\frac{-c}{n_{m\beta}-1}\frac{\partial w_{m\beta}}{\partial p^{*}_{gm\beta}} + \frac{b}{n_{f\beta}}\frac{\partial w_{m\beta}}{\partial p_{gf\beta}}\right)
\end{aligned} \tag{34.3}
$$

$$
\begin{aligned}
F_{mm}^{\beta a} = & -c\frac{\partial w_{m\beta}}{\partial p_{im\beta}} + \frac{b}{n_{m\beta}-1}\frac{\partial w_{m\beta}}{\partial p^{*}_{gm\beta}} + \\
& \left(n_{m\beta}-1\right) r_{mm}^{\beta a}\left[\frac{b}{n_{m\beta}-1}\frac{\partial w_{m\beta}}{\partial p_{im\beta}} + \frac{1}{n_{m\beta}-1}\left[\left(n_{m\beta}-2\right)\frac{b}{n_{m\beta}-1} - c\right]\frac{\partial w_{m\beta}}{\partial p^{*}_{gm\beta}}\right]
\end{aligned} \tag{34.4}
$$

##### 3.4.2 The behaviour with mortality consequences

As for deriving (33.1) – (34.4), the expressions accounting for *mortality* outcomes of the behaviour, expressed by the focal in given *sex* and age $a$ in an $\alpha$-type group, directed to group-mates in given *sex*, are derived as below ('$M$' indicates the behaviour has *mortality* impacts — its first and second subscripts indicate the *recipient* and *focal* sex, respectively):

$$\begin{aligned} M_{ff}^{\alpha a} = & c\frac{\partial w_{f\alpha}}{\partial \mu_{if\alpha}} + \frac{-b}{n_{f\alpha}-1}\frac{\partial w_{f\alpha}}{\partial \mu_{gf\alpha}^{*}} + \\ & \left(n_{f\alpha}-1\right) r_{ff}^{\alpha a}\left[\frac{-b}{n_{f\alpha}-1}\frac{\partial w_{f\alpha}}{\partial \mu_{if\alpha}} + \frac{1}{n_{f\alpha}-1}\left[\left(n_{f\alpha}-2\right)\frac{-b}{n_{f\alpha}-1}+c\right]\frac{\partial w_{f\alpha}}{\partial \mu_{gf\alpha}^{*}}\right] + \\ & n_{m\alpha} r_{mf}^{\alpha a}\frac{c-b}{n_{f\alpha}}\frac{\partial w_{m\alpha}}{\partial \mu_{gf\alpha}} \end{aligned} \tag{35.1}$$

$$\begin{aligned} M_{mf}^{\alpha a} = & c\frac{\partial w_{f\alpha}}{\partial \mu_{if\alpha}} + \frac{-b}{n_{m\alpha}}\frac{\partial w_{f\alpha}}{\partial \mu_{gm\alpha}} + \\ & \left(n_{f\alpha}-1\right) r_{ff}^{\alpha a}\left(\frac{c}{n_{f\alpha}-1}\frac{\partial w_{f\alpha}}{\partial \mu_{gf\alpha}^{*}} + \frac{-b}{n_{m\alpha}}\frac{\partial w_{f\alpha}}{\partial \mu_{gm\alpha}}\right) + \\ & n_{m\alpha} r_{mf}^{\alpha a}\left(\frac{-b}{n_{m\alpha}}\frac{\partial w_{m\alpha}}{\partial \mu_{im\alpha}} + \frac{-b}{n_{m\alpha}}\frac{\partial w_{m\alpha}}{\partial \mu_{gm\alpha}^{*}} + \frac{c}{n_{f\alpha}}\frac{\partial w_{m\alpha}}{\partial \mu_{gf\alpha}}\right) \end{aligned} \tag{35.2}$$

$$\begin{aligned} M_{fm}^{\alpha a} = & c\frac{\partial w_{m\alpha}}{\partial \mu_{im\alpha}} + \frac{-b}{n_{f\alpha}}\frac{\partial w_{m\alpha}}{\partial \mu_{gf\alpha}} + \\ & n_{f\alpha} r_{fm}^{\alpha a}\left(\frac{-b}{n_{f\alpha}}\frac{\partial w_{f\alpha}}{\partial \mu_{if\alpha}} + \frac{-b}{n_{f\alpha}}\frac{\partial w_{f\alpha}}{\partial \mu_{gf\alpha}^{*}} + \frac{c}{n_{m\alpha}}\frac{\partial w_{f\alpha}}{\partial \mu_{gm\alpha}}\right) + \\ & \left(n_{m\alpha}-1\right) r_{mm}^{\alpha a}\left(\frac{c}{n_{m\alpha}-1}\frac{\partial w_{m\alpha}}{\partial \mu_{gm\alpha}^{*}} + \frac{-b}{n_{f\alpha}}\frac{\partial w_{m\alpha}}{\partial \mu_{gf\alpha}}\right) \end{aligned} \tag{35.3}$$

$$\begin{aligned} M_{mm}^{\alpha a} = & c\frac{\partial w_{m\alpha}}{\partial \mu_{im\alpha}} + \frac{-b}{n_{m\alpha}-1}\frac{\partial w_{m\alpha}}{\partial \mu_{gm\alpha}^{*}} + \\ & n_{f\alpha} r_{fm}^{\alpha a}\frac{c-b}{n_{m\alpha}}\frac{\partial w_{f\alpha}}{\partial \mu_{gm\alpha}} + \\ & \left(n_{m\alpha}-1\right) r_{mm}^{\alpha a}\left[\frac{-b}{n_{m\alpha}-1}\frac{\partial w_{m\alpha}}{\partial \mu_{im\alpha}} + \frac{1}{n_{m\alpha}-1}\left[\left(n_{m\alpha}-2\right)\frac{-b}{n_{m\alpha}-1}+c\right]\frac{\partial w_{m\alpha}}{\partial \mu_{gm\alpha}^{*}}\right] \end{aligned} \tag{35.4}$$

In a $\beta$-type group, the corresponding expressions are:

$$M_{ff}^{\beta a} = c\frac{\partial w_{f\beta}}{\partial \mu_{if\beta}} + \frac{-b}{n_{f\beta}-1}\frac{\partial w_{f\beta}}{\partial \mu^*_{gf\beta}} + \left(n_{f\beta}-1\right) r_{ff}^{\beta a}\left[\frac{-b}{n_{f\beta}-1}\frac{\partial w_{f\beta}}{\partial \mu_{if\beta}} + \frac{1}{n_{f\beta}-1}\left[\left(n_{f\beta}-2\right)\frac{-b}{n_{f\beta}-1}+c\right]\frac{\partial w_{f\beta}}{\partial \mu^*_{gf\beta}}\right] + n_{m\beta} r_{mf}^{\beta a}\frac{c-b}{n_{f\beta}}\frac{\partial w_{m\beta}}{\partial \mu_{gf\beta}} \tag{36.1}$$

$$M_{mf}^{\beta a} = c\frac{\partial w_{f\beta}}{\partial \mu_{if\beta}} + \frac{-b}{n_{m\beta}}\frac{\partial w_{f\beta}}{\partial \mu_{gm\beta}} + \left(n_{f\beta}-1\right) r_{ff}^{\beta a}\left(\frac{c}{n_{f\beta}-1}\frac{\partial w_{f\beta}}{\partial \mu^*_{gf\beta}} + \frac{-b}{n_{m\beta}}\frac{\partial w_{f\beta}}{\partial \mu_{gm\beta}}\right) + n_{m\beta} r_{mf}^{\beta a}\left(\frac{-b}{n_{m\beta}}\frac{\partial w_{m\beta}}{\partial \mu_{im\beta}} + \frac{-b}{n_{m\beta}}\frac{\partial w_{m\beta}}{\partial \mu^*_{gm\beta}} + \frac{c}{n_{f\beta}}\frac{\partial w_{m\beta}}{\partial \mu_{gf\beta}}\right) \tag{36.2}$$

$$M_{fm}^{\beta a} = c\frac{\partial w_{m\beta}}{\partial \mu_{im\beta}} + \frac{-b}{n_{f\beta}}\frac{\partial w_{m\beta}}{\partial \mu_{gf\beta}} + n_{f\beta} r_{fm}^{\beta a}\left(\frac{-b}{n_{f\beta}}\frac{\partial w_{f\beta}}{\partial \mu_{if\beta}} + \frac{-b}{n_{f\beta}}\frac{\partial w_{f\beta}}{\partial \mu^*_{gf\beta}} + \frac{c}{n_{m\beta}}\frac{\partial w_{f\beta}}{\partial \mu_{gm\beta}}\right) + \left(n_{m\beta}-1\right) r_{mm}^{\beta a}\left(\frac{c}{n_{m\beta}-1}\frac{\partial w_{m\beta}}{\partial \mu^*_{gm\beta}} + \frac{-b}{n_{f\beta}}\frac{\partial w_{m\beta}}{\partial \mu_{gf\beta}}\right) \tag{36.3}$$

$$M_{mm}^{\beta a} = c\frac{\partial w_{m\beta}}{\partial \mu_{im\beta}} + \frac{-b}{n_{m\beta}-1}\frac{\partial w_{m\beta}}{\partial \mu^*_{gm\beta}} + n_{f\beta} r_{fm}^{\beta a}\frac{c-b}{n_{m\beta}}\frac{\partial w_{f\beta}}{\partial \mu_{gm\beta}} + \left(n_{m\beta}-1\right) r_{mm}^{\beta a}\left[\frac{-b}{n_{m\beta}-1}\frac{\partial w_{m\beta}}{\partial \mu_{im\beta}} + \frac{1}{n_{m\beta}-1}\left[\left(n_{m\beta}-2\right)\frac{-b}{n_{m\beta}-1}+c\right]\frac{\partial w_{m\beta}}{\partial \mu^*_{gm\beta}}\right] \tag{36.4}$$

##### 3.4.3 Inclusive Fitness Impacts of Helping/Harming

Equations (33.1) – (36.4) above explicitly specify whether helping/harming is directed to the female or male groupmates of a focal individual in given sex, age, and grouptype. With these, we can finally derive the *expected* fecundity or mortality impacts of the behaviour on a *random* groupmate of the focal.

To illustrate, for example,

$$\begin{aligned} F_f^{\alpha a} &= \frac{n_{f\alpha}-1}{n_{f\alpha}-1+n_{m\alpha}} \times F_{ff}^{\alpha a} + \frac{n_{m\alpha}}{n_{f\alpha}-1+n_{m\alpha}} \times F_{mf}^{\alpha a} \\ &= \frac{\left[(n_{f\alpha}-1) \times F_{ff}^{\alpha a} + n_{m\alpha} \times F_{mf}^{\alpha a}\right]}{n_{f\alpha}-1+n_{m\alpha}} \end{aligned} \tag{37}$$

captures the weighted average *fecundity* impacts of the behaviour on a random groupmate (either sex) of a focal *female* aged $a$ in an $\alpha$-type group (where $\frac{n_{f\alpha}-1}{n_{f\alpha}-1+n_{m\alpha}}$ or $\frac{n_{m\alpha}}{n_{f\alpha}-1+n_{m\alpha}}$ is the probability that helping/harming is directed to a female or male groupmate, respectively), while, similarly,

$$M_m^{\beta a} = \frac{\left[n_{f\beta} \times M_{fm}^{\beta a} + (n_{m\beta}-1) \times M_{mm}^{\beta a}\right]}{n_{f\beta}+n_{m\beta}-1} \tag{38}$$

captures the expected *mortality* impacts of the behaviour on a random groupmate of a focal *male* aged $a$ in an $\beta$-type group.

Analogous expressions $F_f^{\beta a}$ and $M_m^{\alpha a}$ apply for the cases where the focal *female* of age $a$ resides in a $\beta$-type group and the focal *male* of age $a$ resides in an $\alpha$-type group, respectively, and are derived by applying exactly the same reasoning as in (37) and (38).

##### 3.4.4 Numerical evaluation of sex-, age-, and grouptype-specific selection

To evaluate the *direction* and *strength* of selection on helping or harming (indicated by the *sign* and the *magnitude* of $c/b^*$, respectively), we analyse its expected fecundity or mortality impacts on a random groupmate of a focal expressing the behaviour, in given sex, age, and grouptype, at the resident demographic equilibrium (Taylor and Frank, 1996, Frank, 1998), where all individuals express the same phenotypic value for fecundity or mortality, consistent with standard invasion analysis in kin selection models (Rousset, 2004). At this equilibrium, a focal individual's own fecundity or mortality, its group's 'excluding-self' average, the whole group's average, and the population's average are all identical (only the relative magnitude of small changes in fecundity or mortality matters for selection as the absolute magnitudes is irrelevant). The personal cost of the behaviour is kept very small so that the standard linear approximation holds for tiny evolutionary steps, giving the exact condition for a new behaviour to spread under weak selection (Hamilton, 1964, Gardner et al., 2011). More specifically, we evaluate the expressions (33.1) – (36.4) derived above under the following conditions to derive the critical $c/b^*$ as a measure of selective pressure on the behaviour (see calculation details in the `Python` script supplemented):

$$p_f = p_{if\alpha} = p_{gf\alpha} = p^*_{gf\alpha} = 1$$
$$p_m = p_{im\alpha} = p_{gm\alpha} = p^*_{gm\alpha} = 1$$
$$p_{if\beta} = p_{gf\beta} = p^*_{gf\beta} = 1$$
$$p_{im\beta} = p_{gm\beta} = p^*_{gm\beta} = 1$$

$$\mu_{if\alpha} = \mu_{gf\alpha} = \mu^*_{gf\alpha} = .1$$
$$\mu_{im\alpha} = \mu_{gm\alpha} = \mu^*_{gm\alpha} = .1$$
$$\mu_{if\beta} = \mu_{gf\beta} = \mu^*_{gf\beta} = .1$$
$$\mu_{im\beta} = \mu_{gm\beta} = \mu^*_{gm\beta} = .1$$

$$c = 10^{-5}$$

#### 4 LOCAL VARIATION IN RECRUITING & SIRING PROBABILITIES

##### 4.1 Local Recruiting Probabilities

In our model, a newly recruited breeder (either sex) in a *larger* group is *more* likely to be native, provided that the larger group contains more female breeders. Specifically, when $n_{f\alpha} < n_{f\beta}$, subtracting (18) from (16) gives

$$\pi_{f\alpha} - \pi_{f\beta} = \frac{\left(1 - d_f\right) d_f \bar{n}_f \left(n_{f\alpha} - n_{f\beta}\right)}{\left[\left(1 - d_f\right) n_{f\alpha} + d_f \bar{n}_f\right]\left[\left(1 - d_f\right) n_{f\beta} + d_f \bar{n}_f\right]} \leq 0 \tag{39}$$

Thus, for $0 < d_f < 1$ and $n_{f\alpha} < n_{f\beta}$, we have $\pi_{f\alpha} < \pi_{f\beta}$: a newly recruited *female* breeder in the larger $\beta$-type group has a higher probability of being native. Similarly, because the production of male offspring also depends on the number of female breeders, when $n_{f\alpha} < n_{f\beta}$, subtracting (19) from (17) gives

$$\pi_{m\alpha} - \pi_{m\beta} = \frac{\left(1 - d_m\right) d_m \bar{n}_f \left(n_{f\alpha} - n_{f\beta}\right)}{\left[\left(1 - d_m\right) n_{f\alpha} + d_m \bar{n}_f\right]\left[\left(1 - d_m\right) n_{f\beta} + d_m \bar{n}_f\right]} \leq 0 \tag{40}$$

Thus, for $0 < d_m < 1$ and $n_{f\alpha} < n_{f\beta}$, we have $\pi_{m\alpha} < \pi_{m\beta}$: a newly recruited *male* breeder in the larger $\beta$-type group also has a higher probability of being native. For each sex-specific expression above, equality is reached when dispersal of that sex is absent, when all offspring of that sex disperse, or when the two grouptypes contain the same number of female breeders. In these cases, group size does not generate difference in the native origin of new recruitments.

##### 4.2 Local Siring Probabilities

In the model, offspring born in a *larger* group are also *more* likely to be sired by a local male (provided that the larger group contains more male breeders). Specifically, when $n_{m\alpha} < n_{m\beta}$, subtracting (21) from (20) gives

$$m_{\alpha} - m_{\beta} = \frac{\left(1 - m\right) \bar{n}_m m \left(n_{m\alpha} - n_{m\beta}\right)}{\left[m n_{m\alpha} + \left(1 - m\right) \bar{n}_m\right]\left[m n_{m\beta} + \left(1 - m\right) \bar{n}_m\right]} \leq 0 \tag{41}$$

Thus, for $0 < m < 1$ and $n_{m\alpha} < n_{m\beta}$, we have $m_{\alpha} < m_{\beta}$: an offspring born in the larger $\beta$-type group has a higher probability of being sired by a local male. Equality is reached when local paternity is absent ($m = 0$), when all offspring are sired locally ($m = 1$), or when the two grouptypes contain the same number of male breeders. In these cases, group size does not generate difference in local siring probability.

#### 5 COMPARISON WITH HOMOGENEOUS POPULATIONS

Comparing distinct, homogeneous populations, each of which contains only the same $\alpha$-type or the same $\beta$-type groups as in the heterogeneous population, across the whale, typical mammal, and ape cases (characterized by clear-cut patterns of local mating and sex-specific dispersal), we find that individuals are more related to their groupmates in smaller groups (Fig. A1), and consequently their social behaviours (helping/harming) are under stronger selective pressures in smaller groups (Fig. A2).

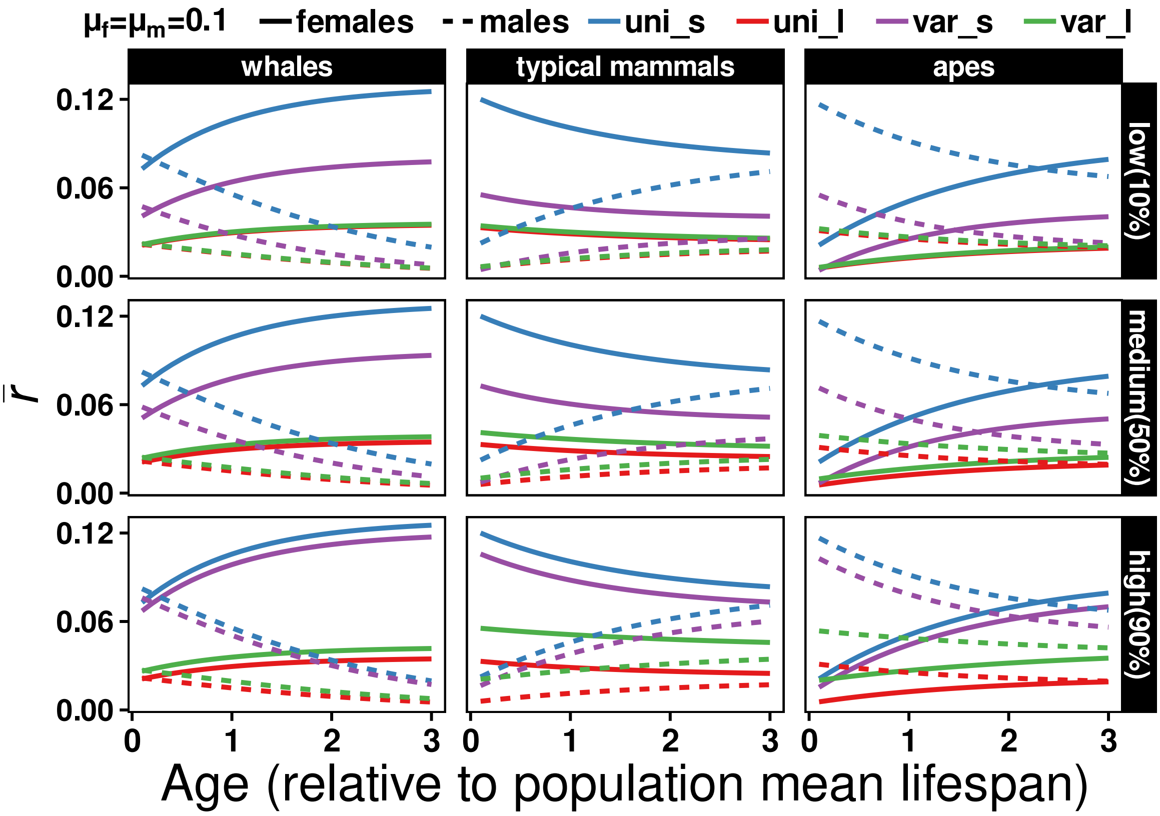

**Fig. A1** Age-specific average relatedness of a female (solid lines) or male (dashed lines) breeder to its groupmates in the two counterpart 'homogeneous' populations made up with either the $\alpha$-type (i.e., smaller groups with 5 females and 5 males, indicated by 'uni_s') or the $\beta$-type (i.e., larger groups with 20 females and 20 males, indicated by 'uni_l') groups — as defined in the 'heterogeneous' population in the main texts (where $\alpha$-type and $\beta$-type groups are indicated by 'var_s' and 'var_l', respectively), across different proportions of smaller groups in the heterogeneous population (low: $u = 10\%$, medium: $u = 50\%$, high: $u = 90\%$).

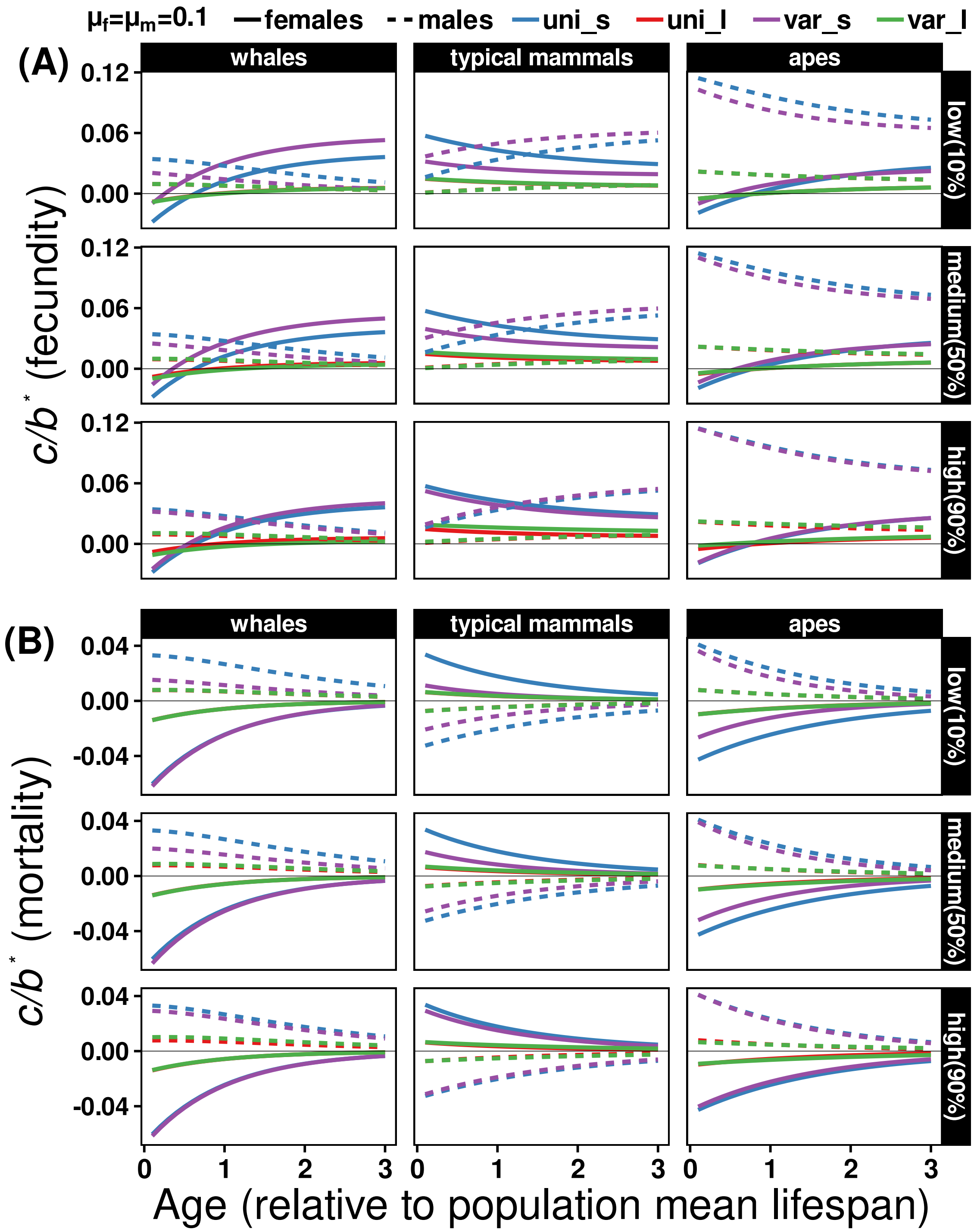

**Fig. A2** Age-specific selective pressures on helping or harming expressed by a female (solid lines) or male (dashed lines) breeder that has fecundity (A) or mortality (B) impact on the individual and its groupmates in the counterpart 'homogeneous' populations made up with only the $\alpha$-type (i.e., smaller groups with 5 females and 5 males, indicated by 'uni_s') or the $\beta$-type (i.e., larger groups with 20 females and 20 males, indicated by 'uni_l') groups — as defined in the 'heterogeneous' population in the main texts (where $\alpha$-type and $\beta$-type groups are indicated by 'var_s' and 'var_l', respectively), across different proportions of smaller groups in the heterogeneous population (low: $u = 10\%$, medium: $u = 50\%$, high: $u = 90\%$). The horizontal lines denote the behaviour has no net fitness impact (selectively neutral).

#### 6 GROUP-SIZE-SPECIFIC RELATEDNESS CANNOT BE INTERPOLATED

While comparisons between distinct, homogeneous populations (Fig. A1) show that smaller groups tend to exhibit higher age-specific relatedness (and stronger selection on helping/harming — Fig. A2), across the *whale*, *ape*, and *typical mammal* cases, this does *not* imply that, under group size local variation, an individuals average relatedness to its groupmates can be simply 'interpolated' from predictions derived for counterpart group-size homogeneous populations in earlier models of kinship dynamics (*nor* does it imply that higher relatedness and stronger selection universally occur in smaller groups — see Fig. A4 and Fig. A5). Rather, in the presence of group size local variation *within* a population, age-specific relatedness in a local group *depends on* the demographic compositions of the *entire* population, and *cannot* be predicted by simple 'interpolation between such counterpart homogeneous populations (Fig. A3).

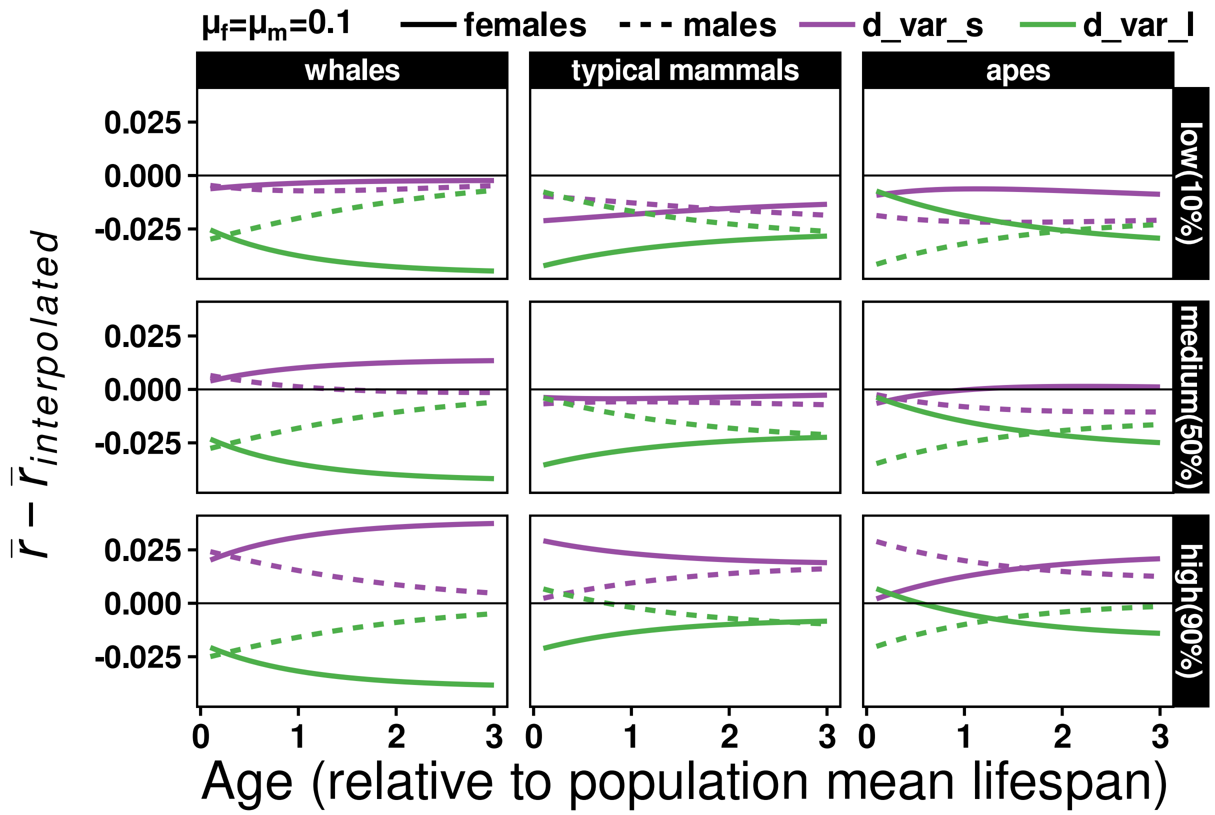

**Fig. A3** An individuals sex-age-specific average relatedness to its groupmates in a heterogeneous population (containing both $\alpha$- and $\beta$-type groups) *cannot* be predicted (or inferred) by simple 'interpolation' from those in two counterpart homogeneous populations (containing only the same $\alpha$-type or the same $\beta$-type groups). The y-axis shows the *relative difference* between sex-age-specific average relatedness in an $\alpha$- ('d_var_s') or $\beta$-type ('d_var_l') group drawn from a heterogeneous population and the corresponding sex-age-specific arithmetic mean derived from two counterpart homogeneous populations — as a simple 'interpolation', across different proportions of smaller ($\alpha$-type) groups in the heterogeneous population (low: $u = 10\%$, medium: $u = 50\%$, high: $u = 90\%$; the horizontal lines denote no relative difference; the $\alpha$- and $\beta$-type groups are the same as those in Fig. A1).

### 7 EXPLORING DEMOGRAPHIC EFFECTS ON KINSHIP DYNAMICS UNDER GROUP SIZE LOCAL VARIATION

In the presence of group size local variation within a population, how do other demographic parameters influence an individual's age-specific relatedness to its groupmates? To place the effects of group size local variation in broader demographic contexts, we conducted a structured comparative exploration of the model's parameter space, systematically varying standard demographic components around a single baseline scenario.

#### 7.1 Hypothetical Demographic Parameter Sets

Specifically, we examined variation in (A) the magnitude of size contrast between smaller and larger groups, (B) sex ratio, implemented as differences in group sex composition, (C) sex-specific mortality rates, (D) the prevalence of local versus non-local mating, (E) sex-specific dispersal rates, and (F) the proportion of smaller versus larger groups in the population (Table A1). Starting from a demographically neutral baseline with balanced sex ratios ($n_{f\alpha} = n_{m\alpha}$, $n_{f\beta} = n_{m\beta}$) in both grouptypes, symmetric mortality and dispersal between the sexes ($\mu_f = \mu_m = .5$, $d_f = d_m = .5$), intermediate local mating ($m = .5$), equal frequencies of smaller and larger groups ($u = .5$), and pronounced local group-size variation (10 vs. 40 individuals), we varied one demographic component at a time while holding all others constant (Table A1). This design allows the effects of each demographic dimension on age-specific relatedness to be directly compared under the same analytical framework.

**Table A1 Hypothetical demographic parameter sets used for comparative exploration of an individual's age-specific relatedness to its groupmates.** Here, a series of targeted demographic perturbations from a neutral reference scenario (R0) are explored. In each block (A–F), only the parameters highlighted relative to the baseline are modified (others are held constant). The $\alpha$ and $\beta$ grouptypes differ in total size, allowing within-population variation in group size and sex composition. Arrows indicate whether a parameter value is increased (↑) or decreased (↓) relative to the baseline. Such a design thus evaluates separately the effects of group-size heterogeneity (A), sex composition (B), mortality rate (C), local mating rate (D), dispersal rate (E), and the proportion of small groups (F) on age-specific relatedness.

| Case | No. | $n_{f\alpha}$ | $n_{m\alpha}$ | $n_{f\beta}$ | $n_{m\beta}$ | $\mu_f$ | $\mu_m$ | $m$ | $d_f$ | $d_m$ | $u$ | Description |
|---|---|---|---|---|---|---|---|---|---|---|---|---|
| R | 0 | 5 | 5 | 20 | 20 | .5 | .5 | .5 | .5 | .5 | .5 | **Baseline**: reference |
| A | 1 | 15↑ | 15↑ | 20 | 20 | .5 | .5 | .5 | .5 | .5 | .5 | **Weaker heterogeneity**: increased $\alpha$ group size |
| | 2 | 5 | 5 | 10↓ | 10↓ | .5 | .5 | .5 | .5 | .5 | .5 | **Weaker heterogeneity**: reduced $\beta$ group size |
| B | 1 | 8↑ | 2↓ | 20 | 20 | .5 | .5 | .5 | .5 | .5 | .5 | **Sex ratio**: ♀-biased in $\alpha$ groups |
| | 2 | 2↓ | 8↑ | 20 | 20 | .5 | .5 | .5 | .5 | .5 | .5 | **Sex ratio**: ♂-biased in $\alpha$ groups |
| | 3 | 5 | 5 | 32↑ | 8↓ | .5 | .5 | .5 | .5 | .5 | .5 | **Sex ratio**: ♀-biased in $\beta$ groups |
| | 4 | 5 | 5 | 8↓ | 32↑ | .5 | .5 | .5 | .5 | .5 | .5 | **Sex ratio**: ♂-biased in $\beta$ groups |
| | 5 | 8↑ | 2↓ | 32↑ | 8↓ | .5 | .5 | .5 | .5 | .5 | .5 | **Sex ratio**: ♀-biased in $\alpha$ & $\beta$ groups |
| | 6 | 8↑ | 2↓ | 8↓ | 32↑ | .5 | .5 | .5 | .5 | .5 | .5 | **Sex ratio**: reverses ♀ bias in $\beta$ groups |
| | 7 | 2↓ | 8↑ | 8↓ | 32↑ | .5 | .5 | .5 | .5 | .5 | .5 | **Sex ratio**: ♂-biased in $\alpha$ & $\beta$ groups |
| | 8 | 2↓ | 8↑ | 32↑ | 8↓ | .5 | .5 | .5 | .5 | .5 | .5 | **Sex ratio**: reverses ♂ bias in $\beta$ groups |
| C | 1 | 5 | 5 | 20 | 20 | .8↑ | .5 | .5 | .5 | .5 | .5 | **Mortality rate**: increases in ♀ |
| | 2 | 5 | 5 | 20 | 20 | .5 | .8↑ | .5 | .5 | .5 | .5 | **Mortality rate**: increases in ♂ |
| | 3 | 5 | 5 | 20 | 20 | .2↓ | .2↓ | .5 | .5 | .5 | .5 | **Mortality rate**: decreases in both sexes |
| | 4 | 5 | 5 | 20 | 20 | .8↑ | .2↓ | .5 | .5 | .5 | .5 | **Mortality rate**: increases in ♀ & decreases in ♂ |
| | 5 | 5 | 5 | 20 | 20 | .2↓ | .8↑ | .5 | .5 | .5 | .5 | **Mortality rate**: decreases in ♀ & increases in ♂ |
| | 6 | 5 | 5 | 20 | 20 | .8↑ | .8↑ | .5 | .5 | .5 | .5 | **Mortality rate**: increases in both sexes |
| D | 1 | 5 | 5 | 20 | 20 | .5 | .5 | .2↓ | .5 | .5 | .5 | **Local mating rate**: decreases |
| | 2 | 5 | 5 | 20 | 20 | .5 | .5 | .8↑ | .5 | .5 | .5 | **Local mating rate**: increases |
| E | 1 | 5 | 5 | 20 | 20 | .5 | .5 | .5 | .8↑ | .5 | .5 | **Dispersal rate**: increases in ♀ |
| | 2 | 5 | 5 | 20 | 20 | .5 | .5 | .5 | .5 | .8↑ | .5 | **Dispersal rate**: increases in ♂ |
| | 3 | 5 | 5 | 20 | 20 | .5 | .5 | .5 | .2↓ | .2↓ | .5 | **Dispersal rate**: decreases in both sexes |
| | 4 | 5 | 5 | 20 | 20 | .5 | .5 | .5 | .8↑ | .2↓ | .5 | **Dispersal rate**: increases in ♀ & decreases in ♂ |
| | 5 | 5 | 5 | 20 | 20 | .5 | .5 | .5 | .2↓ | .8↑ | .5 | **Dispersal rate**: decreases in ♀ & increases in ♂ |
| | 6 | 5 | 5 | 20 | 20 | .5 | .5 | .5 | .8↑ | .8↑ | .5 | **Dispersal rate**: increases in both sexes |
| F | 1 | 5 | 5 | 20 | 20 | .5 | .5 | .5 | .5 | .5 | .1↓ | **% small groups**: reduced |
| | 2 | 5 | 5 | 20 | 20 | .5 | .5 | .5 | .5 | .5 | .9↑ | **% small groups**: increased |

##### 7.2 Effects of Standard Demographic Paramters

The results of our exploration on kinship dynamics (Fig. A4) are briefly summarized as below.

**A. Group-size heterogeneity:**

Lower group size heterogeneity reduces between-grouptype difference in the probability that two individuals sampled from the same group share a parent. Enlarging smaller groups (A1) reduces the probability that two individuals in smaller groups share recent ancestors (i.e., weaker genetic drift, see 1.3), whereas shrinking larger groups (A2) increases the probability of shared ancestry among individuals in larger groups (i.e., stronger drift); as a result, an individual's average relatedness to others changes in a more similar pattern between smaller and larger groups, compressing between-grouptype differences in patterns of kinship dynamics.

**B. Sex composition (breeder sex ratio):**

Sex-ratio skew affects kinship dynamics by shaping the relative contribution of maternal versus paternal lineages to local ancestry — it does not uniformly strengthen or weaken kinship ties, but affects how ancestry is distributed between female and male lineages. Reducing the number of breeders of one sex elevates age-specific relatedness in that sex, while typically depressing it in the more common sex (B1–B4, B5–B8). That is, when one sex is rarer, ancestry through that sex is drawn from fewer parents, increasing the probability that individuals share ancestry through that sex (i.e., stronger drift) and thereby elevating relatedness for individuals of the rarer sex.

**C. Mortality rate:**

Mortality affects how fast breeders are replaced by new recruits (recall that at each timestep breeding vacancies left by deaths are fully claimed by offspring winning the fair-lottery competition), thereby shaping how fast an individual's relatedness to others changes with age. Specifically, with higher mortality rate (e.g., C6), individuals are replaced more frequently and their kinship ties change faster with age (indicated by the increased skewness in the distributions of age-specific relatedness in C6), while udner lower mortality (e.g., C3), individuals persist longer and their kinship ties change slower with age (indicated by the decreased skewness in the distributions of age-specific relatedness in C3).

**D. Local mating rate:**

By regulating the retention versus replacement of local ancestry, local mating affects the overall level of age-specific relatedness across all sex-grouptype classes. Increasing local mating strengthens relatedness by increasing the probability that offspring share local ancestry (D2), whereas reducing it weakens local relatedness due to higher probability of introducing non-local lineages (D1). These effects are stronger in the male than female sex, as non-local mating introduces unrelated paternal rather than maternal lineages (even when individuals remain in their natal group).

**E. Dispersal rate:**

Dispersal impacts kinship dynamics by introducing non-local lineages. Reduced dispersal preserves local ancestry, whereas increased dispersal introduces immigrant lineages that dilute local relatedness, modulating age-specific relatedness coherently across all sex-grouptype classes (e.g., E3, E6). Sex-biased dispersal (E4–E5) shapes how ancestry is partitioned maternally and paternally, creating sex-specific responses in kinship dynamics. Besides, sex-biased dispersal affects relatedness in both sexes, as ancestry is jointly inherited through the female and male lineages; increasing dispersal in one sex dilutes ancestry through that sex for all in a group, thereby shaping age-specific relatedness even among individuals of the less-dispersing sex (i.e., spillover effect, E1 & E2).

**F. Population composition:**

When smaller groups are more common, lineages are more likely to pass through these groups characterized by stronger genetic drift, where ancestry concentrates more rapidly, elevating age-specific relatedness across sex-grouptype classes (F2), whereas when larger groups dominate, ancestry is more diffuse, reducing age-specific relatedness (F1).

Collectively, these results show that group size local variation sets the basic demographic context in which other standard demographic processes shape the patterns of kinship dynamics in local groups. Importantly, under such within-population variation, the age-specific relatedness of a focal individual in a given grouptype cannot be inferred from that group's own demographies alone: demographic conditions in other grouptype also modulate the ancestry contributing to the focal grouptype, thereby modulating the emergent patterns of kinship dynamics therein.

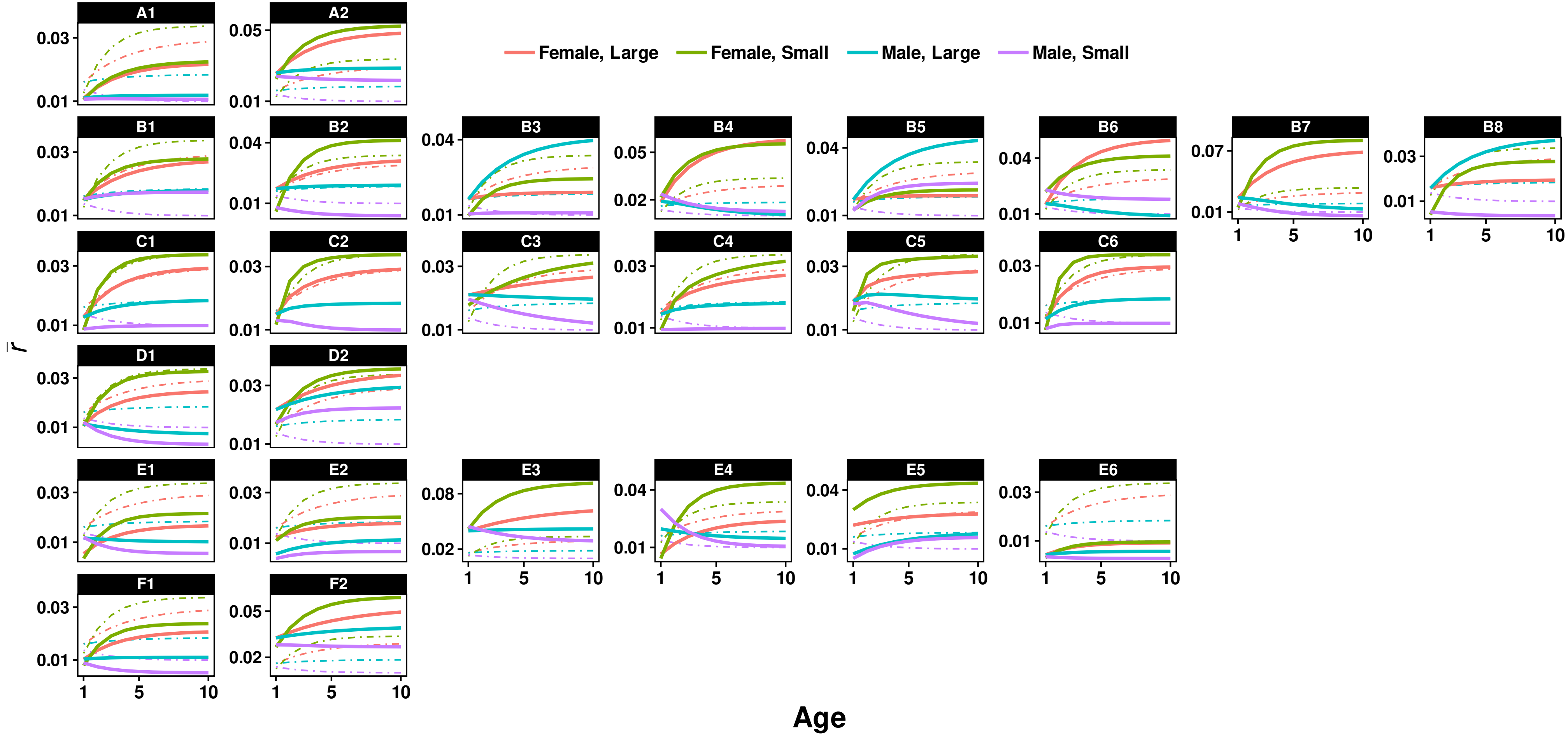

**Fig. A4 Kinship dynamics across sex-grouptype classes under group size local variation and different demographic regimes.** Model parameters were perturbed around a neutral baseline (R0; Table A1) to examine the effects of (A) group-size heterogeneity, (B) sex ratios, (C) sex-specific mortality rates, (D) local mating rate, (E) sex-specific dispersal rates, and (F) grouptype composition on kinship dynamics in populations with group size local variation. Solid lines show the age-specific average relatedness of a focal individual to its groupmates (for clarity up to age 10, after which they approach their asymptotes) in each sex-grouptype class (shown with colors) under each demographic regime (A1–F2 in Table A1), while dot-dashed lines show the corresponding sex-grouptype-specific patterns under the baseline demographic regime (R0).

### 8 SEX DIFFERENCES IN KINSHIP DYNAMICS

#### 8.1 Identifying Demographic Drivers

Here, we identify the demographic processes that generate sex differences in kinship dynamics. For simplicity, we generate predictions using the parameter sets listed in Table A1, but assume *completely* local mating (i.e., $m = 100\%$) across all cases (A – F) — *except* for the reference case R0, where local versus non-local mating rates remain balanced (i.e., $m = 50\%$).

Collectively, the results of our exploration (Fig. A5) show that sex differences in the patterns of kinship dynamics emerge from extra-group mating (D1 or D2 versus R0), as well as sex biases in breeder number (B1 – B8), mortality (C1 – C2 & C4 – C5), and dispersal (E1 – E2 & E4 – E5), and are independent of group size heterogeneity (A1 – A2) and grouptype composition within a population (F1 – F2).

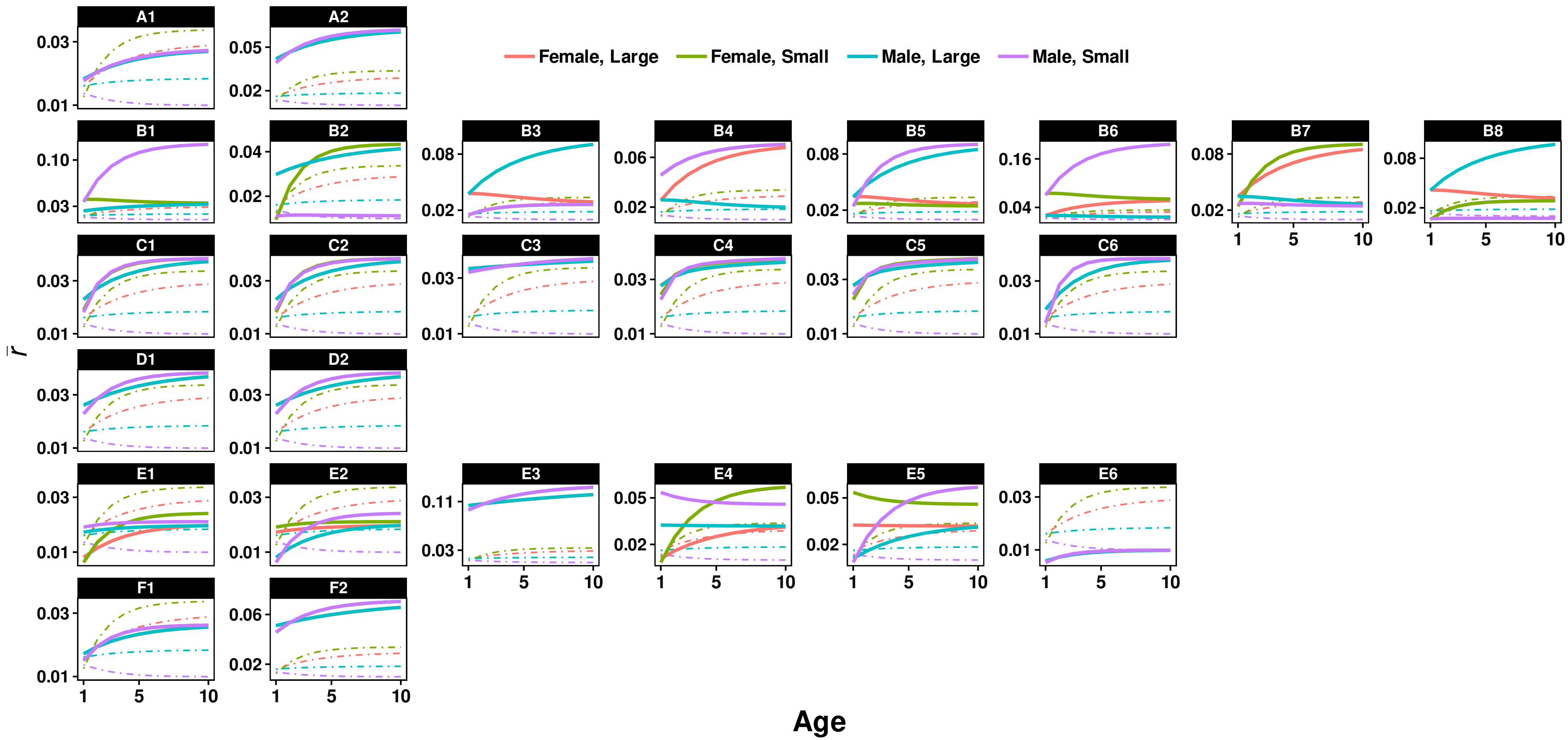

**Fig. A5 Sex differences in kinship dynamics emerge from extra-group mating, and sex biases in breeder number (sex ratio), mortality rate, and dispersal rate.** Here, solid lines across cases A – F show the age-specific average relatedness of a focal individual to its groupmates (for clarity up to age 10, after which they approach their asymptotes) in each sex-grouptype class (shown with colors) under each demographic regime realized by the parameter values in Table A1 — *except* assuming *completely* local mating (i.e., $m = 100\%$); dot-dashed lines show the corresponding sex-grouptype-specific patterns of kinship dynamics under the baseline demographic regime (i.e., R0 in Table A1, where $m$ remains at 50%), which are the same across the panels.

#### 8.2 Exploring The Role of Dispersal and Mating

As shown above, dispersal and local mating are key determinants of kinship dynamics: dispersal *dilutes* local relatedness, whereas local mating *enhances* it (Section 7), and sex differences in kinship dynamics can arise from sexbiased dispersal and extra-group mating (Section 8.1). However, these results do not clarify how dispersal and local mating jointly shape sex differences in kinship dynamics. Here, we address this gap in the presence and absence of group size local variation. Specifically, we explore how sex difference are shaped by (I) female and male dispersal (under completely local mating), (II) female dispersal and local mating (under moderate male dispersal), and (III) male dispersal and local mating (under moderate female dispersal). For clarity, predictions are generated by varying the focal parameters (dispersal and/or local mating rates) around the reference R0 (Table A1), while all other parameters remain consistent — for homogeneous populations (without group size local variation), the universal group sizes (small or large) correspond to those in R0 for heterogeneous populations. We quantify the magnitude of sex differences with the root mean square (RMS) of age-specific differences in average local relatedness between the sexes.

We found that when mating is entirely restricted to local groups ($m = 1$), increasing sex bias in dispersal leads to larger sex differences in kinship dynamics (these differences diminish as dispersal becomes more balanced between the sexes). For a given degree of dispersal bias, sex differences are more pronounced in smaller groups, both in the presence (Fig. A6-A) and absence (Fig. A7-A) of group size local variation. When male/female dispersal is fixed at a moderate level ($d_m = .5$ or $d_f = .5$), the effect of female/male dispersal on sex differences in kinship dynamics depends on the prevalence of local mating (and vice versa). Such interdependence is stronger in smaller groups and this holds regardless of if group size local variation is present (Fig. A6, B & C) or not (Fig. A7, B & C). Moreover, when one sex is with moderate dispersal rate, lower local mating rate and/or lower dispersal rate in the other sex are associated with larger sex differences in kinship dynamics, while moderate dispersal rates in both sexes, combined with completely local mating, tend to minimize the magnitude of sex differences in kinship dynamics (Fig. A6, B & C; Fig. A7, B & C).

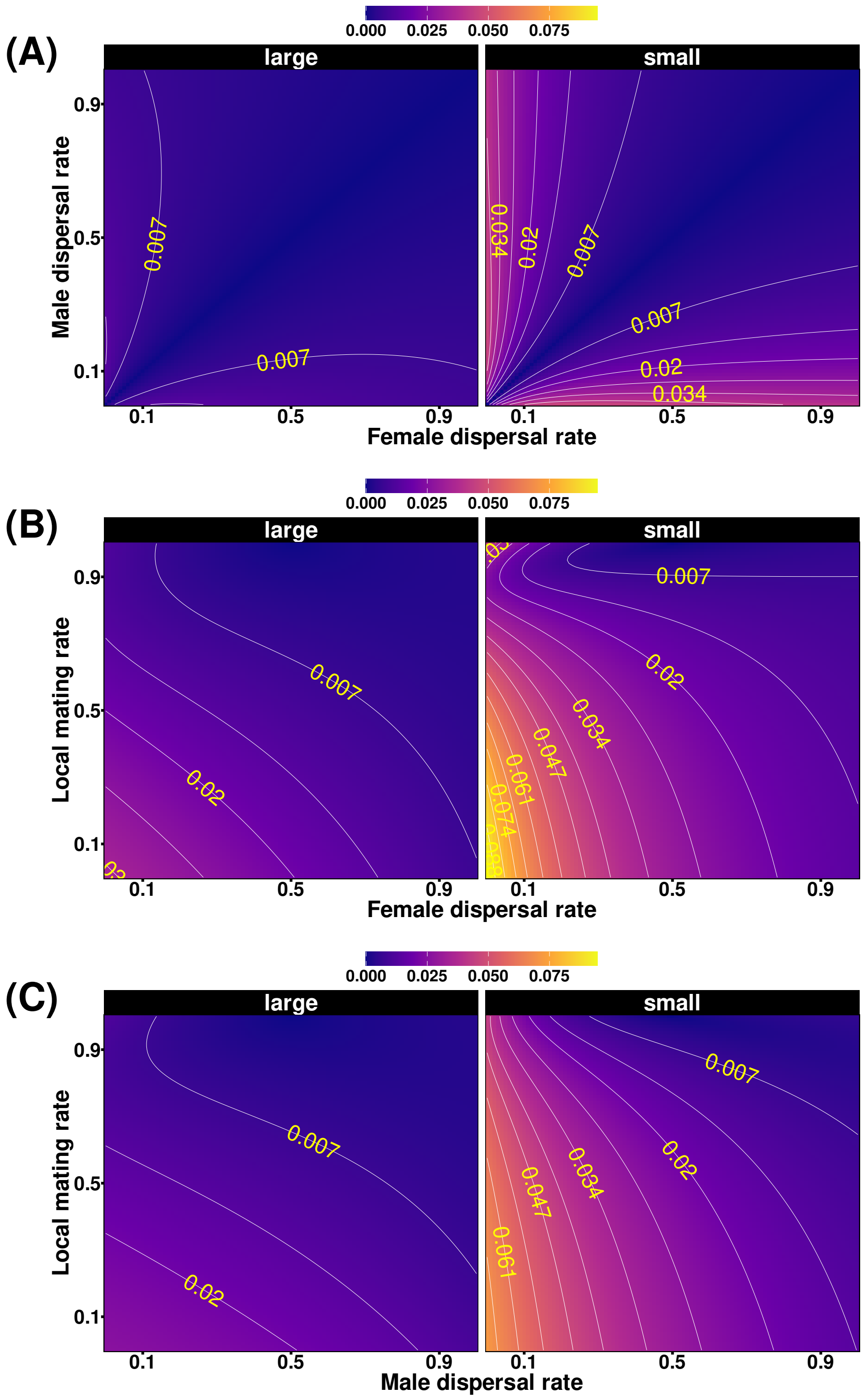

**Fig. A6. The role of dispersal and local mating in shaping sex differences in kinship dynamics in the *presence* of group size local variation within a population.** Sex differences in kinship dynamics are quantified as the root mean square (RMS, shown by colour scale) of age-specific differences in average local relatedness between the sexes in coexisting smaller and larger groups within a population. Panels show the joint effects of (A) female and male dispersal under completely local mating ($m = 1$), (B) female dispersal and local mating under moderate male dispersal ($d_m = .5$), and (C) male dispersal and local mating under moderate female dispersal ($d_f = .5$) on RMS. Contour lines connect parameter combinations (dispersal and/or local mating rates on the axes) that generate the same magnitude of RMS (values shown on the lines). All other parameter values remain as those in R0 (Table A1).

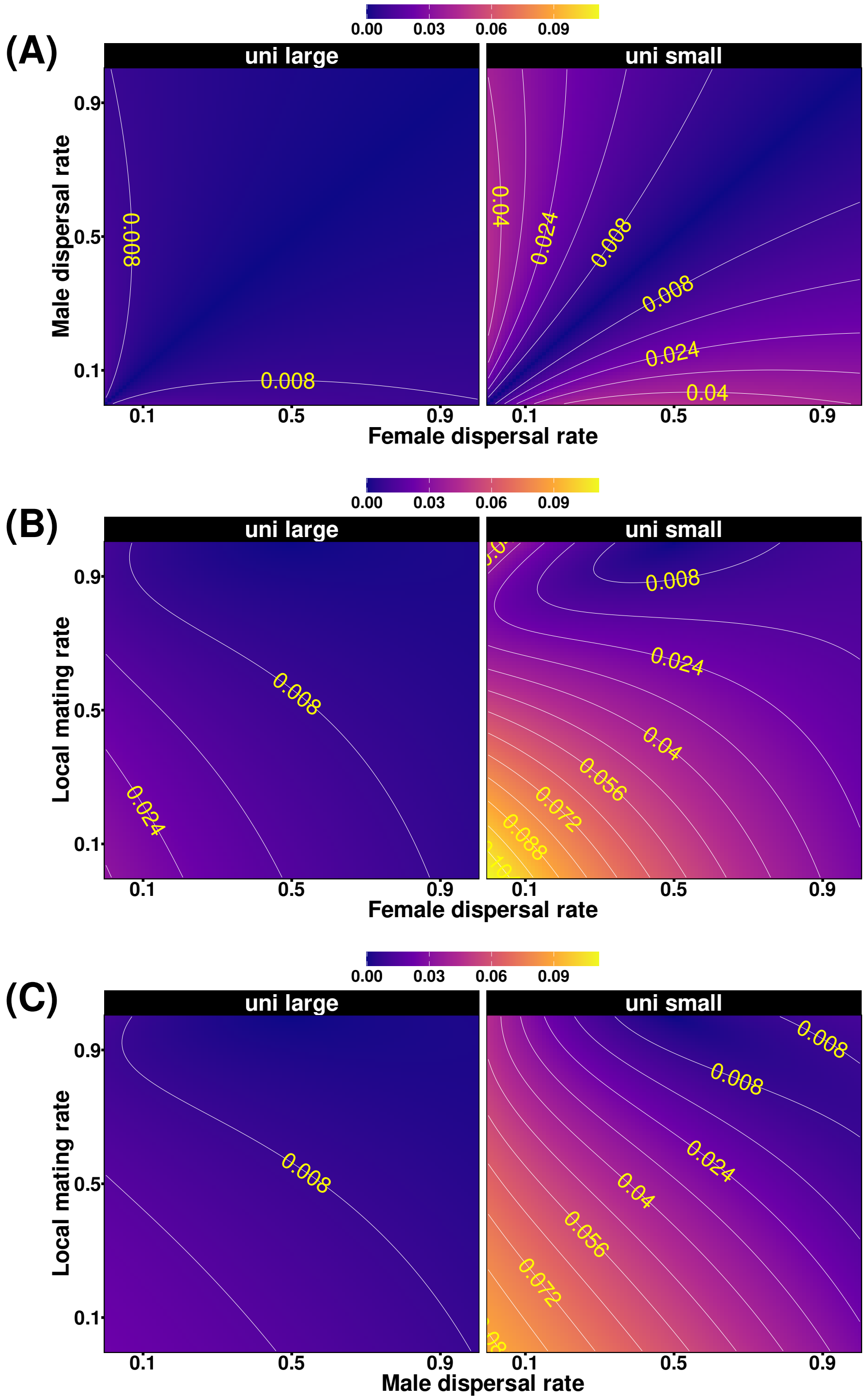

**Fig. A7. The role of dispersal and local mating in shaping sex differences in kinship dynamics in the *absence* of group size local variation (i.e., in homogeneous populations).** Sex differences in kinship dynamics are quantified as the root mean square (RMS, shown by colour scale) of age-specific differences in average local relatedness between the sexes in two distinct group-size homogeneous populations. Panels show the joint effects of (A) female and male dispersal under completely local mating ($m = 1$), (B) female dispersal and local mating under moderate male dispersal ($d_m = .5$), and (C) male dispersal and local mating under moderate female dispersal ($d_f = .5$) on RMS. In this context, the two universal group sizes ('uni small' and 'uni large') are the same as those specified in R0 for the group-size heterogeneous population, but here represent two distinct, group-size homogeneous populations. Contour lines connect parameter combinations (dispersal and/or local mating rates on the axes) that generate the same magnitude of RMS (values shown on the lines). All other parameter values remain as those in R0 (Table A1).

#### 9 FEMALE KILLER WHALE REPRODUCTIVE RATE

Here, we illustrate the uncertainty associated with empirically testing our predictions on grouptype-linked variations in social life history traits. In this example, we specifically focus on examining how the reproductive rates of younger versus older female southern resident killer whales (*Orcinus orca*) vary across social groups of different sizes, using a long-term (1976 – 2022) dataset maintained by the Center for Whale Research, comprising annual records of both social group sizes and females' birth occurrences across these groups, collected through complete individual-level censuses using photo-identification methods (Bigg et al., 1990) under research permits issued by the USA (NMFS 27308) and Canada (DFO-SARA 288).

Individuals (of both sexes) in this population form a multilevel society characterized by fission-fusion dynamics but preferentially associate into three pods, namely the $K$, $J$, and $L$ pod (Bigg et al., 1990, Ellis et al., 2022). In this analysis, we define these pods as social groups, and consider only those reproductive females which aged between 9 and 45 years old (inclusive). As a result, the $K$ pod is identified as the smallest group (on average 17.19 reproductive females in each census year), while the $J$ pod as the medium-sized group (on average 20.91 reproductive females in each census year) and the $L$ pod as the largest group (on average 43.47 reproductive females in each census year; Fig. A8-A). Females in the reproductive age window across these pods are classified as either $young$ or $old$ age class, depending on whether they are below the median age of reproductive females (i.e., 24) in this population.

We analysed annual birth records using a binomial generalized linear mixed model (GLMM) with a logit link, treating birth occurrence as a Bernoulli response, female identity as a random intercept (to account for repeated observations in the census dataset), and age class, pod identity, and their interaction as fixed effects (see R script supplemented for details). The likelihood-ratio test comparing the fitted model to a random-intercept-only reference model indicated *non-significant* statistical association between female birth probability in a census year and age class, pod identity, or their interaction ($\chi^2 = 5.57, df = 5, p = .35$), despite variations in the *GLMM-estimated* birth rates (Fig. A8-B) are qualitatively consistent with our evolutionary predictions.

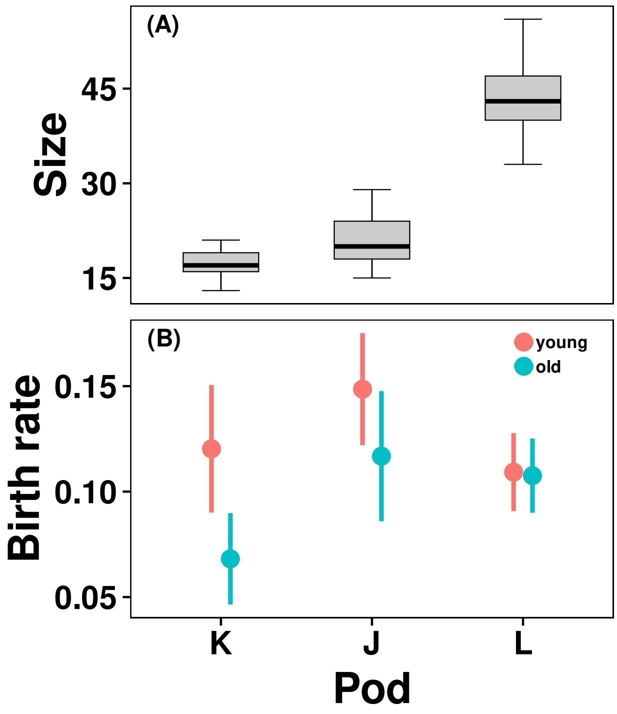

**Fig. A8.** Variation in pod size (A) and model-estimated annual birth probability of young ($9 \leq \text{age} < 24$) versus old ($24 \leq \text{age} \leq 45$) female southern resident killer whales across the *K*, *J*, and *L* pods (B). Pod-size distributions are derived from complete annual census data collected between 1976 and 2022. Birth probabilities were estimated from a binomial generalized linear mixed model with a logit link and female identity as a random effect, fitted to individual-level female birth records (age 24 corresponds to the median reproductive age and age 45 to the average age at last reproduction). Points denote model estimates and vertical bars indicate standard errors (young: red; old: blue). Likelihood-ratio test indicated statistically non-significant associations between birth probability and age class, pod identity, or their interaction — here estimates are presented to visualise the magnitude and uncertainty of plausible differences given the data.
